# Impairment of autophagy-lysosomal activity near the chronically implanted microelectrodes

**DOI:** 10.1101/2023.05.31.543108

**Authors:** Keying Chen, Camila Garcia Padilla, Kirill Kiselyov, Takashi Kozai

## Abstract

Intracortical microelectrodes that can record and stimulate brain activity have become a valuable technique for basic science research and clinical applications. However, long-term implantation of these microelectrodes can lead to progressive neurodegeneration in the surrounding microenvironment, characterized by elevation in disease-associated markers. Dysregulation of autophagy-lysosomal degradation, a major intracellular waste removal process, is considered a key factor in the onset and progression of neurodegenerative diseases. It is plausible that similar dysfunctions in autophagy-lysosomal degradation contribute to tissue degeneration following implantation-induced focal brain injury, ultimately impacting recording performance. To understand how the focal, persistent brain injury caused by long-term microelectrode implantation impairs the autophagy-lysosomal pathway, we employed two-photon microscopy and immunohistology. This investigation focused on the spatiotemporal characterization of autophagy-lysosomal activity near the chronically implanted microelectrode. We observed an aberrant accumulation of immature autophagy vesicles near the microelectrode over the chronic implantation period. Additionally, we found deficits in autophagy-lysosomal clearance proximal to the chronic implant, which was associated with an accumulation of autophagy cargo and a reduction in lysosomal protease level during the chronic period. Furthermore, our evidence suggests astrocytes contribute to the clearance of myelin debris via autophagy-lysosomal degradation near the implanted microelectrode. Together, this study sheds light on the process of brain tissue degeneration caused by long-term microelectrode implantation, with a specific focus on impaired intracellular waste degradation.

## 1. Introduction

Intracortical microelectrodes, offering high-resolution neural monitoring and modulation, hold promise for research and clinical use. [1–5]. However, their long-term performance and potential tissue damage remain crucial factors that need to be addressed to meet the clinical requirements for device application throughout a patient’s lifetime. Studies have shown a decline in long-term microelectrode functional performance in rodents and non-human primates, eventually resulting in device failure and loss of neural signal detection [6–8]. The recording performance decline coincides with neuronal loss near long-term microelectrodes and the accumulation of disease-linked factors like hyperphosphorylated tau. [9]. Deficits in intracellular machinery for waste removal, such as autophagy-lysosomal degradation, have recently been associated with the build-up of hyperphosphorylated tau in neurodegenerative diseases. [10, 11]. Yet, it is unknown how this autophagy-lysosomal pathway is disrupted by the long-term microelectrode implantation as well as its relationship to chronic tissue degeneration.

Studies show that microelectrode implantation results in acute neuroinflammation (0-2 weeks post-implantation) followed by chronic foreign body responses (4+ week implantation) [12, 13]. Acute neuroinflammation is initiated by the insertion of a stiff microelectrode, rupturing the vascular network [14, 15]. The infiltrated blood cells and plasma proteins trigger microglia, astrocytes, and NG2-glia to become activated with a polarized morphology, extend toward the implant within the first week post-implantation, and form an encapsulating sheath of gliosis near the microelectrode by two weeks [16–19]. Microglia and astrocytes during this acute neuroinflammatory phase take on an important phagocytic capacity to remove damaged cells and proteins [20, 21]. This phagocytic activity is mediated, at least in part, by autophagy-lysosome activity, an intracellular organelle that plays important roles in regulating cellular survival, morphologies, and functions [22, 23].

Additionally, autophagosomes andlysosomes play crucial roles in neuronal degeneration and progressive demyelination, which primarily occur during the later chronic foreign body response phase [12, 24–27]. Oligodendrocytes and myelin provide axons with electrical insulation and metabolic support [28]. Given the autophagy-lysosomal-endosome axis’s role in oligodendrocyte differentiation and myelin growth, the failure of oligodendrocyte regeneration and myelin density loss near long-term microelectrodes imply potential dysregulation of autophagy-lysosome activity. [29, 30]. For neurons, intracellular autophagy-lysosome trafficking is involved in maintaining cell survival, mitochondria integrity, and synaptic plasticity, and thus is critical for neuronal health and action potential generation [31]. However, the decreased neuronal density and increased neuronal apoptosis at the chronic implantation period suggest neuronal autophagy-lysosomal dysfunction [30, 32, 33]. Together, these cellular changes in brain tissue near the microelectrode point to autophagy lysosomal activity as a potential key regulator that impacts both the neuronal activity and glial scarring.

The autophagy-lysosomal pathway is an intracellular process crucial for cell health and activity. [23, 34, 35]. In this process, the cargo material is engulfed by a membrane sack that then seals and forms a closed, double-layer organelle known as the autophagosome [36]. Next, the autophagosomes are fused with lysosomes that contain proteases and hydrolases to form autolysosomes for the cargo degradation [37]. The lysosomal proteases and hydrolases that degrade the waste products are activated at acidic pH. This acidification, maintained by V-ATPase proton pumps, is an ATP consuming, metabolically costly process. [38]. The autophagy-lysosome pathway is crucial for timely self-degradation of misfolded protein aggregates and damaged cytoplasmic organelles, including mitochondria. [39–41]. Accumulated misfolded protein aggregates are toxic and this accumulation is a hallmark of many neurodegenerative pathologies [42, 43]. Furthermore, the chronic presence of dysfunctional mitochondria can lead to excessive generation of reactive oxygen species (ROS) [44]. Recent observations show that the autophagy-lysosomal pathway is involved in neurogenesis, synapse plasticity, and cellular remodeling, underscoring its importance for brain tissue integrity [45, 46].

However, deficits in the autophagy-lysosomal pathway are often observed at the onset and during the progression of neurodegenerative diseases [47–49]. In an Alzheimer’s disease (AD) mouse model, excessive accumulation of autophagy vacuoles containing amyloid β was observed in dystrophic neurites [50]. Additionally, genetic mutations of proteins involved in the autophagy-lysosomal pathway cause neurodegenerative syndromes [51–54]. Although brain trauma shares similar pathologies and exhibits an accumulation of neurodegenerative disease-associated factors [55, 56], it is unknown whether and how autophagy-lysosomal activity is impaired by focal, long-term brain injury caused by microelectrode implantation. Filling this knowledge gap helps advance our current understanding of chronic tissue degeneration near the implanted microelectrode and may identify novel pathways for improving neural implant biocompatibility.

To investigate spatiotemporal changes in autophagy-lysosomal activity near the chronically implanted microelectrodes, we applied two-photon imaging of the CAG-LC3b-RFP-EGFP mouse model. This transgenic mouse line has a dual fluorescent, pH-sensitive sensor on LC3B-binding autophagy vesicles, allowing monitoring of the real-time autophagy lysosomal dysregulation near the microelectrode over time [57–59]. We also used immunohistology to study the pathologies of the autophagy-lysosome pathway over 84 days of chronic microelectrode implantation. Here, we hypothesized that the device implantation injury dysregulates autophagy and lysosomal activity at the chronic microelectrode interface, resulting in deficits in lysosomal degradation capability and, thus, the accumulation of cargo containing vesicles. *In vivo* imaging demonstrated an abnormal accumulation of immature autophagy vesicles at chronic 63-77 days post-implantation. Immunohistological results demonstrated that reactive astrocytes increased the lysosomal population proximal to the microelectrode at the chronic implantation period. Additionally, there was a discrepancy between the elevated lysosome population and the reduced level of a lysosomal protease, suggesting an impaired autophagy-lysosomal degradation capability. These results highlight a novel autophagy-lysosome activity perspective on chronic tissue health near the chronically implanted microelectrodes that point to new targets along the autophagy lysosome axis for promoting long-term device performances.

## 2. Methods

### 2.1. Microelectrode angled implantation surgery

The CAG-LC3b-RFP-EGFP transgenic mouse line (C57BL/6-Tg[CAG-RFP/GFP/Map1lc3b]1Hill/J, Jackson Laboratories; Bar Harbor, ME) was used (n = 5, 8-week male, 25–30 g) to track real-time autophagy degradation [57–59]. Mice received bilateral craniotomy windows and a 4-shank Michigan-style non-functional silicon microelectrode (A4x4-3mm-100–703 CM16; NeuroNexus, Ann Arbor, MI), as described previously [60]. Briefly, mice were first anesthetized by an intraperitoneal (IP) injection of 75 mg/kg ketamine and 7 mg/kg xylazine. Ketamine update (40 mg/kg) was used to maintain anesthesia during surgery. Then mice were head-fixed in a stereotaxic frame on a heating pad and administered with continuous supply of oxygen (1 L/min). Two stainless-steel bone screws were implanted over motor cortices for headcap stability. Next, bilateral craniotomies (3 x 3 mm) were drilled over visual cortex in coordinates that 1 mm anterior to lambda and 1.5 mm lateral from the midline. Saline was periodically applied to maintain brain hydration and avoid thermal damage. The non-functional microelectrode was inserted 600 μm into the cortex at a 30° angle (insertion velocity of 400 μm/s, oil hydraulic Microdrive; MO-82, Narishige, Japan), avoiding large vessels with tip depth of ∼ 300 μm below the brain surface [15]. The craniotomy windows were sealed with Kwik-sil sealant (World Precision Instruments, Sarasota County, FL) and then sealed with glass coverslips and dental cement [61]. All experimental procedures were conducted following approval by the University of Pittsburgh, Division of Laboratory Animal Resources, and Institutional Animal Care and Use Committee in accordance with the standards for humane animal care as set by the Animal Welfare Act and the National Institutes of Health Guide for the Care and Use of Laboratory Animals.

### 2.2. Two-photon chronic imaging

Data acquisition by two-photon imaging was illustrated in Fig. 1A. Mice were situated on a 4-wheel rotating treadmill for awake, head-fixed imaging. The imaging setup consisted of a two-photon laser scanning microscope with an OPO laser and second fixed 1040 nm laser (Insight DS+; Spectra-Physics, Menlo Park, CA), non-descanned photomultiplier tubes (Hamamatsu Photonics KK, Hamamatsu, Shizuoka, Japan), and a 16x, 0.8 numerical aperture water immersion objective lens (Nikon Instruments, Melville, NY). CAG-LC3b-RFP-EGFP mice were imaged using dual excitation lasers at 920 nm and 1040 nm for optimal excitation of EGFP and RFP. The total laser power was kept below 40 mW throughout the imaging sessions. Z-stacks were acquired every 2 µm along the microelectrode shank to the tip. An equivalent control tissue volume was imaged on the no-implant contralateral side. Each image in z-stacks had a resolution of 1024 × 1024 pixels (550.5 × 550.5 μm). Daily imaging was performed until day 14, followed by weekly imaging up to day 84 post implantation.

**Figure 1.**
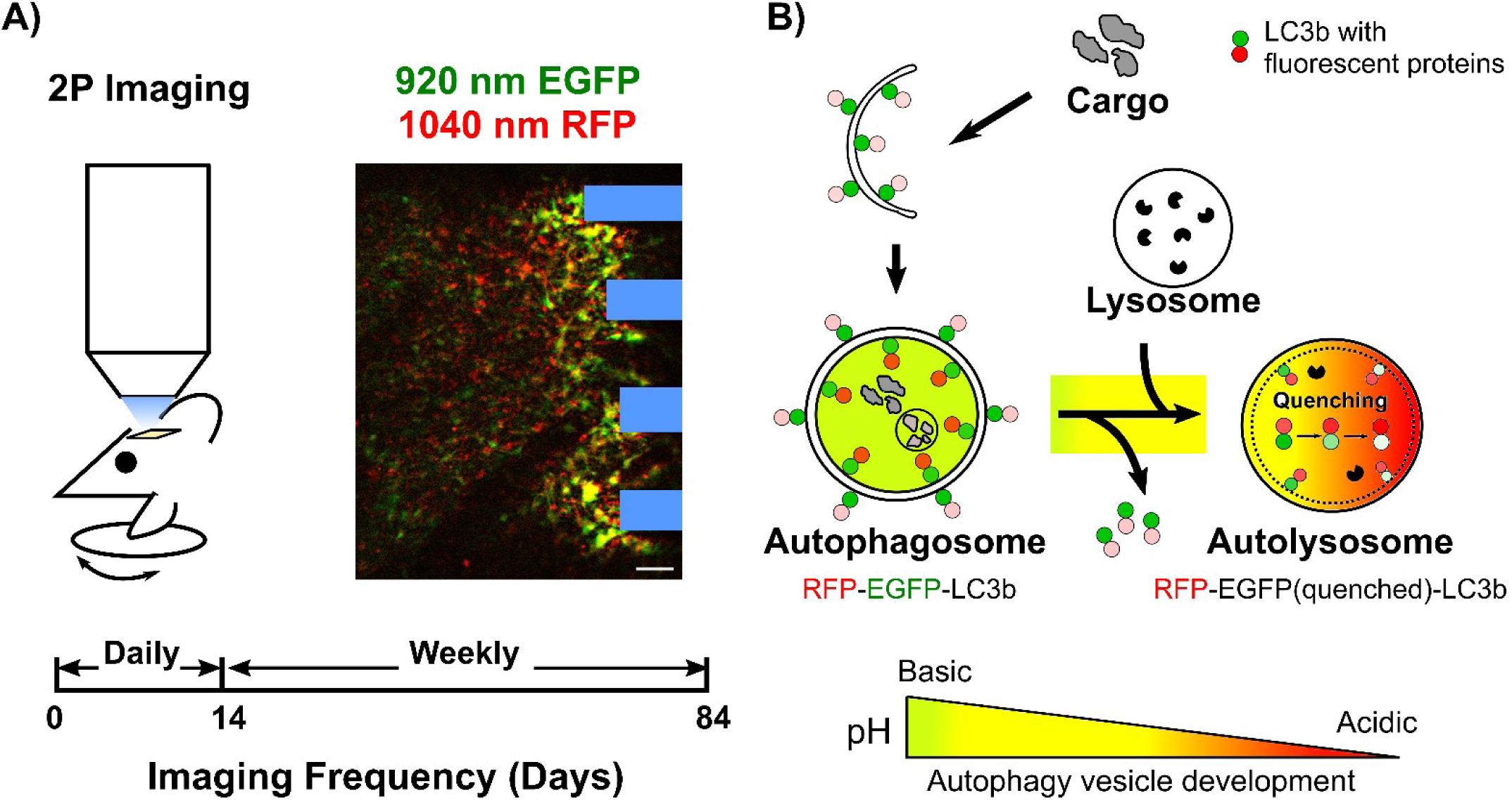
Two-photon imaging of intracellular autophagy activity in the dual fluorescent CAG-LC3b-RFP-EGFP mice. **A)** Awake transgenic mice with cranial windows were head-fixed on a rotating treadmill imaged using a two-photon microscope. Dual lasers at 920 nm and 1040 nm wavelengths were used for simultaneous RFP and EGFP excitation *in vivo*. Laser power was kept below 40 mW to avoid thermal damage. Imaging frequency was daily in the first two weeks post-implantation and then weekly until week 12 post-implantation. The implanted 4-shank microelectrode is highlighted in blue. Scale bar = 50 μm. **B)** Schematic representation of the autophagy-lysosome pathway with dual fluorescence-tagged LC3b. The tagged LC3b consist of pH-resistant RFP and pH-sensitive EGFP, enabling tracking of autophagy dynamics. Autophagy involves the engulfment of cargo by a membrane sack, forming autophagosomes. Autophagosomes then fuse with lysosomes to form autolysosomes for cargo degradation. Immature autophagy vesicles express both EGFP and RFP, while mature autophagy vesicles that have undergone acidification only express RFP. The transition from green/yellow to red signals indicates vesicle acidification.

### 2.3 Immunohistochemistry and Confocal Imaging

Single shank non-functional Michigan-style microelectrodes (A16-3 mm-50-703-CM16; NeuroNexus, Ann Arbor, MI) were perpendicularly implanted into the left visual cortex of C57BL6 wildtype mice [33]. The surgical preparations followed the angled implantation procedure described above, except that the microelectrode shank was perpendicularly inserted at a speed of 15 mm/s using a DC motor-controller (C-863, Physik Instructmente, Karlsruhe, Germany). The shank was implanted 1,600 µm into the brain. Mice were sacrificed at days 7, 14, 28, and 84 post-implantation, with 4 mice for each time point. Transcardial perfusion with 1x PBS and then 4% paraformaldehyde (PFA) was performed on the mice. The extracted brains were then post-fixed in 4% PFA at 4°C for 18 hours. Then the extracted brains were rehydrated in 30% sucrose and embedded into optimum cutting temperature (OCT) media. These brains were sectioned horizontally into 10 µm slices from the surface of the brain to a depth of 1,600 µm. For staining, tissue sections first underwent heat-induced antigen retrieval (0.1 M citric Acid, 0.1 M sodium citrate) and endogenous peroxidase blocking. Then sections were permeabilized and blocked by 0.1% Triton-X with 10% normal goat serum in PBS at room temperature for 1 h. Next, tissue sections were blocked by the endogenous mouse immunoglobulin G (IgG) with donkey anti-mouse IgG fragment (Fab) for 2 hours at 1:10 dilution. Primary antibodies were applied overnight at 4°C. All primary antibodies used in this study are listed in Table 1. In the following day, tissue sections were first washed with 1x PBS and then incubated with secondary antibodies of donkey anti-rat Alexa Fluor 405 (1:500, Abcam, ab175670), donkey anti-rabbit Alexa Fluor 488 (1:500, Abcam, ab150061), donkey anti-rat Alexa Fluor 488 (1:500, Abcam, ab150153), donkey anti-mouse 568 (1:500, Abcam, ab175700), donkey anti-goat 647 (1:500, Abcam, ab150135), donkey anti-chicken 647 (1:500, JacksonImmunoResearch, 730-605-155) in the dark for 2 hours. Sections were washed and mounted with Fluoromount G media (SouthernBiotech, #0100-20) and glass covers.

| Primary Antibody | Target | Supplier | Host | Dilution |
| --- | --- | --- | --- | --- |
| Anti-LC3B | Autophagy vesicles | Abcam (ab51520) | Rabbit | 1:1000 |
| Anti- NeuN | Neurons | Sigma (MAB2300) | Mouse | 1:500 |
| Anti- IBA1 | Microglia | Millipore (MABN92) | Mouse | 1:500 |
| Anti-GFAP | Astrocytes | Abcam (ab4674) | Chicken | 1:500 |
| Anti-CC1 | Oligodendrocyte soma | Millipore (OP80) | Mouse | 1:100 |
| Anti-MBP | Myelin | Abcam (ab7349) | Rat | 1:500 |
| Anti-LAMP1 | Lysosome vesicles | Abcam (ab25245) | Rat | 1:500 |
| Anti-TFEB | Lysosome biogenesis | Sigma (SAB4503154) | Rabbit | 1:500 |
| Anti-PLP | Myelin | Sigma (SAB2101830) | Rabbit | 1:500 |
| Anti-SQSTM1/p62 | Autophagy clearance | Abcam (ab91526) | Rabbit | 1:500 |
| Anti-Cathepsin D | Lysosomal protease | Abcam (ab6313) | Mouse | 1:100 |
| Anti-TOMM20 | Mitochondria | Abcam (ab186735) | Rabbit | 1:100 |
| Anti-4-Hydroxynonenal | Oxidative Stress | Abcam (ab48506) | Mouse | 1:50 |

Each section was imaged individually with a confocal microscope (FluoView 1000, Olympus, Inc., Tokyo, Japan) equipped with a 20x, oil-immersive objective lens. TIFF images were captured with the electrode track centered, in a resolution of 16 bit, 1024 × 1024 pixels (635.9 × 635.9 μm).

### 2.4. Data analysis

#### 2.4.1 Two photon data processing

The z-stacks acquired by two-photon imaging of CAG-LC3b-RFP-EGFP transgenic mice were used to dynamically track the real-time autophagy-lysosome activity near the microelectrode throughout the 84-day post-implantation period. Due to potential variations in imaging parameters and craniotomy window quality across time points, intensity-based analysis was deemed unsuitable for quantifying autophagy-lysosome activity. Instead, the density and morphologies of EGFP and RFP activity were analyzed to investigate the temporal changes in autophagy activity.

To mitigate interference from meningeal tissue, each z-stack was rotated using ImageJ’s “Interactive Stack Rotation” until the dura surface appeared flat. The total tissue volume below the dura was then summed using the z-projection function. Then background noise was corrected using the background subtraction method with a rolling ball radius of 10 pixels on the original sum z-projected images (**Supplementary Fig. 1A-B**). This correction method effectively reduced uneven background noise while preserving the fluorescent signal. Next, Ostu’s thresholding was applied to generate a binary mask of fluorescent signal (**Supplementary Fig. 1C**). To validate that the thresholding algorithm did not unevenly filter out fluorescent signals, the histogram distributions of mean-normalized intensity of filtered background pixels were compared between implant and contralateral windows (**Supplementary Fig. 1D**). Probability density functions of these distributions were plotted (**Supplementary Fig. 1E**). No significant difference was detected in the background pixel intensity distributions between implant and contralateral sides (Two-sample Kolmogorov–Smirnov test, p = 0.996). This noise minimization procedure was performed on each raw two-photon stacks prior to density and morphology analysis.

The density of autophagy vesicles was quantified by calculating the pixel area percentage of fluorescent signals (RFP+, EGFP+, RFP+EGFP-) within the region of interest (ROI). ROIs chosen at the 4-shank microelectrode interface were rectangular measuring 270 μm × 480 μm (500 pixels x 900 pixels) and centered between the second and third shanks of the microelectrode. Corresponding ROIs of the same size were chosen in the contralateral hemisphere. The densities of RFP+ signals, EGFP+ signals, RFP+ EGFP signals, and the ratio of EGFP+ density over RFP+ density were measured in both the implant and contralateral tissue at each time point.

Fluorescent clusters larger than 4 pixel^2^ (1 μm^2^) in the RFP and EGFP channels were identified using the “Analyze particle” ImageJ plugin. Cluster density was calculated as the total number of clusters divided by the ROI area in mm^2^. The ROIs used for cluster density analysis were the same ROIs used for pixel density quantification. Additionally, the mean sizes of RFP and EGFP clusters were recorded at each time point. Changes in cluster density and mean size of RFP and EGFP clusters on the implant and contralateral sides were plotted over time.

To assess autophagy activity near the electrode-tissue interface, RFP and EGFP activity over the implant surface was measured. Each raw z-stack was rotated until the probe surface was horizontal to the XY plane. The tissue volume within 20 μm above the implant surface was summed via z-projection. Background subtraction and Otsu’s thresholding were applied to generate a binary mask. The implant surface was outlined as the ROI, and the surface coverage was calculated as fraction of fluorescent signal (RFP+, EGFP+, RFP+ EGFP) over the ROI. Furthermore, mean sizes of RFP and EGFP clusters within probe surface ROI were tracked over time.

To investigate the spatial distributions of autophagy activity near the microelectrode, RFP and EGFP clusters within 300 μm of the outmost microelectrode shank were quantified. The distance of each cluster from the microelectrode shank was calculated based on its XY coordinates. RFP and EGFP clusters were counted and plotted over time and over distance as spatiotemporal heatmaps. Additionally, changes in clusters counts over time were plotted for each spatial bin as an alternative visualization method.

#### 2.4.2. Histological data processing

The previously published MATLAB script, I.N.T.E.N.S.I.T.Y. was used to process the fluorescent intensity as previously described [62, 63]. The center of the implant hole was manually identified, and spatial bins were automatically generated at 10 µm intervals up to 300 µm from the probe. The average grayscale intensity within each spatial bin was calculated by taking the mean value of pixels above the thresholding (1.5 standard deviations above background noise). Fluorescence intensities were then normalized to the average intensity from the four corners of each image (greater than 300 μm away from the implant hole). The normalized intensity was averaged across all animals at each time point and plotted over distances. Bar graphs generated to display the averaged intensity within 50 µm spatial bins up to 150 µm from the implant site.

Colocalization analyses were performed by merging the autophagy marker (LC3B) or lysosome marker (LAMP1) with various cellular markers (NeuN/IBA1/GFAP/CC1/MBP/PLP) using ImageJ. Binary masks of each fluorescent marker were generated using background subtraction (rolling ball radius = 10 pixels) followed by Ostu’s thresholding algorithm. The colocalization percentage of LC3B or LAMP1 with specific cellular markers was determined by calculating the proportion of overlapping pixels (nonzero in both binary masks) relative to the total number of nonzero pixels in the LC3B or LAMP1 binary mask. For cell counting analysis, neurons with Transcription factor EB (TFEB) nuclear translocation were identified as DAPI+ TFEB+ NeuN+ cells in spatial bins spaced 50 µm apart up to 300 µm from the implant site. The density of DAPI+ TFEB+ NeuN+ cells was calculated by dividing the total cell counts by the tissue area per spatial bin. Data was averaged across all animals at each time point and plotted over distances.

### 2.5. Statistics

Two-way ANOVA with Tukey post-hoc (p < 0.05) was used to analyze the two-photon data, assessing significant differences in autophagy RFP and EGFP pixel density, cluster size, cluster density, and surface coverages between day 0 and other time points. One-way ANOVA followed by Dunnett’s test (p < 0.05) was employed to identify significant differences in clusters at each spatial bin and metrics. Immunohistological data were analyzed using a two-way ANOVA with Tukey post-hoc (p < 0.05) to determine significant differences in normalized intensity, colocalization percentage, and cell density between different time points of implantation. Bonferroni correction was applied for multiple comparisons as appropriate.

## 3. Results

Given the crucial role of autophagy-lysosomal activity in tissue homeostasis and neuroinflammation response [64, 65], we aimed to reveal spatiotemporal changes in autophagy-lysosomal activity near the implanted microelectrode. To monitor real-time autophagy degradation in response to microelectrode implantation over time *in vivo*, we used the CAG-LC3b RFP-EGFP transgenic mice (**Fig. 1A**). This transgenic model allows the tagged LC3B protein to express dual fluorescence, RFP and EGFP [57–59]. The tandem fluorescent LC3B binds to autophagosome membrane and persists during autophagosome development and even after lysosomal fusion, until they are fully degraded with the internal cargo by the lysosomal proteases [66]. The EGFP (pKa = 5.9) is sensitive to acidic environment and thus undergoes fluorescent quenching after lysosomal fusion [67]. Instead, the RFP (pKa = 4-5) is relatively acid-insensitive [57]. Thus, the inclusion of both RFP and EGFP in this transgenic model allows the observation of dynamic autophagy degradation, which depends on the acidity of the environment (**Fig 1B**). Our dual laser wavelength setting allowed the optimal excitation of both EGFP (920 nm) and RFP (1040 nm), as the total laser power was less than 40 mW to avoid thermal damage.

The dual fluorescence of EGFP and RFP serves as an indicator of pH levels within LC3B-tagged autophagy vesicles, representing the population of autophagosomes [11]. As autophagosomes fuse with lysosomes and become autolysosomes, the internal environment inside autolysosomes is gradually acidified. This acidification leads to the quenching of EGFP fluorescence causing a shift in color from yellow to orange and then to red [11]. The expression of RFP eventually decreases after autophagy clearance of tandem fluorescent LC3B. With this LC3B model, we were able to track the activity of immature autophagosomes (yellow/green) and mature, acidified autolysosomes (orange/red) near the microelectrode throughout the implantation period.

### 3.1. Aberrant accumulation of immature autophagy vesicles around the chronically implanted microelectrodes

The implantation of microelectrode elicits a series of biological responses, including acute neuroinflammation (0-2 weeks post-implantation) and chronic foreign body responses (4+ weeks post-implantation). Brain cells demonstrate distinct patterns of morphologies, functionality, and survival during these two temporal phases. Autophagy activity dynamically responds to stimuli resulting in increased autophagosome formation, lysosomal fusion, and autolysosome population [68]. Therefore, we asked how the autophagy activity changes near the implanted microelectrode over time. The representative images showed that the autophagy activity at the electrode-tissue interface was low on the implantation surgery day (day 0) but increased over the first week during the early chronic period, and then persisted at an elevated level in the area adjacent to the chronic implant (**Fig. 2A**). Thus, we first aimed to investigate the time course of population changes of total autophagy (RFP+) vehicles, autophagosome (EGFP+RFP+), and autolysosomes (EGFP-RFP+) around chronically implanted microelectrodes.

**Figure 2.**
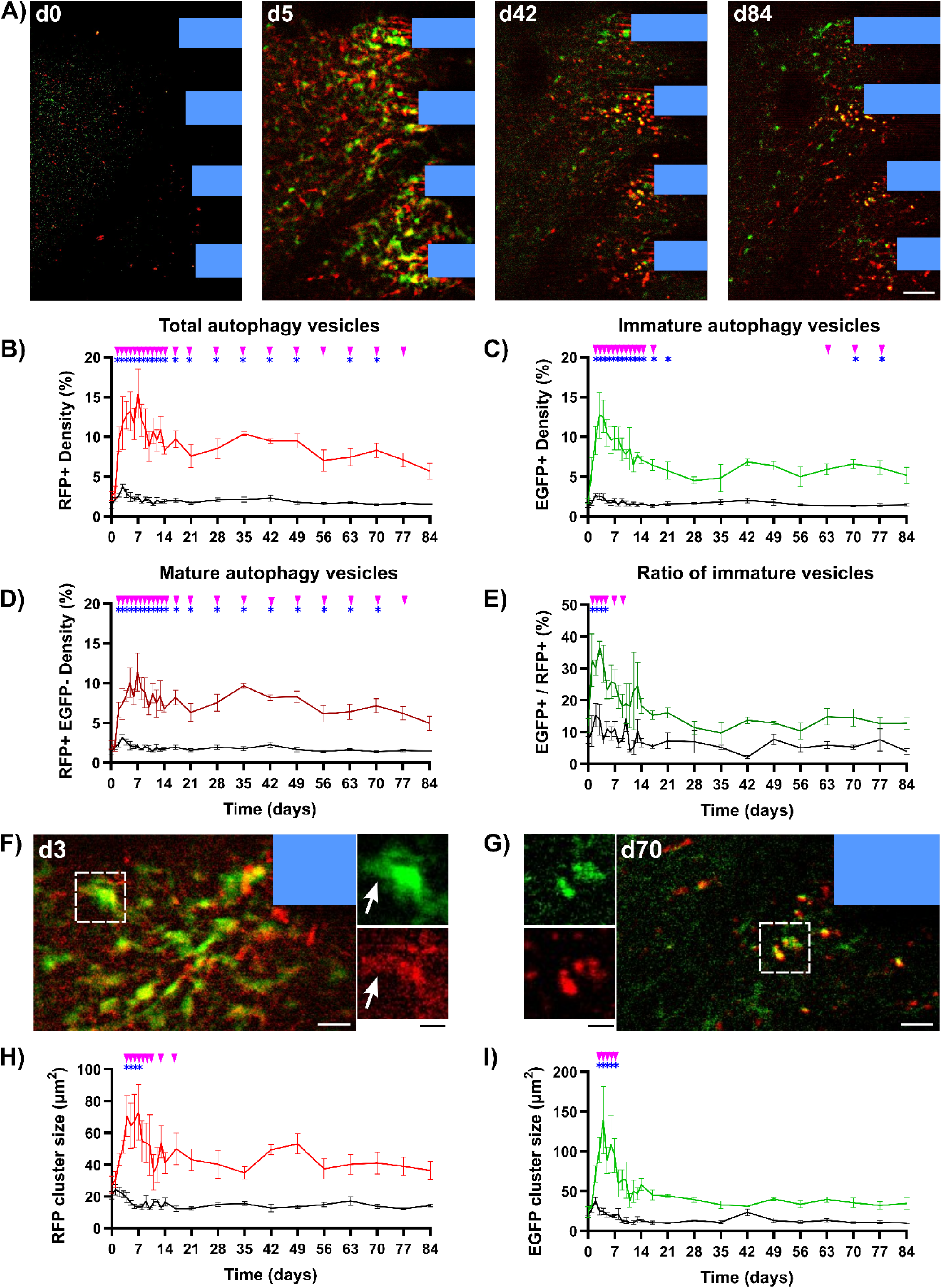
Chronic accumulation of immature, partially acidified autophagy vesicles following microelectrode implantation. **A)** Representative two-photon images showing autophagy granules surrounding the implanted 4-shank microelectrode array (highlighted in blue) over 84-days post-implantation in the CAG-LC3b-RFP-EGFP mice. Scale bar = 50 μm. **B)** Quantification of RFP+ pixel density at microelectrode interfaces (colored) and no-implant contralateral side (black) over time. RPF+ pixel density, representing the total autophagy vesicle population, was significantly higher near the implant compared to the contralateral control through day 84 post-implantation. **C)** EGFP pixel density, indicative of immature autophagy vesicle (autophagosomes), exhibited a transient peak within 21-days post-implantation and chronic accumulation during days 63 - 77 post-implantation. **D)** The pixel density of RFP+ EGFP-, representing acidified autophagy vesicles (autolysosomes), remained persistently higher near the microelectrode interfaces throughout the implantation period. **E)** The ratio of EGFP pixel density to RFP pixel density was significantly higher compared to the contralateral control within 14-days post-implantation, indicating an acute predominance of immature autophagy vesicles (autophagosomes) near the implanted microelectrode. **F)** Representative two-photon images of dual-fluorescent labeled autophagy cluster adjacent to the implanted microelectrode (blue) on day 3 post-implantation. White scale bar = 20 μm. Magnified images (on the right) show a large bright body and shaded, glial like processes of an autophagy cluster. Black scale bar = 10 μm. **G)** Representative two-photon images of small, dot-like morphologies of an autophagy cluster near the probe on day 70 post-implantation. Change in cluster sizes of RFP **(H)** and EGFP **(I)** signals over the 84-day implantation period between the implant and contralateral ROI regions in the CAG-LC3b-RFP-EGFP mice. Significant increases in EGFP cluster sizes occurred at days 3 – 7 post-implantation, while significant increases in RFP cluster sizes appeared at days 4 – 17 post-implantation compared to the contralateral sides. Significance between implant and contralateral sides at each time point was marked in pink solid triangles (Two-way ANOVA with Tukey post-hoc, *p* < 0.05). Significant comparisons between day 0 and other time points at implant ROIs were labeled as blue asterisks (two-way ANOVA with Tukey post-hoc, *p* < 0.05). No significance was found between time points at contralateral regions. All data were presented as mean ± standard deviation.

To quantitatively examine the changes in the total population of autophagy vesicles near the implanted microelectrode, pixel density of pH-resistant RFP was measured at the implant side and contralateral (craniotomy window only) side over time (**Fig. 2B**). RFP pixel density in the implant side peaked at day 7 and was significantly higher than contralateral tissue during days 2-77 post-implantation (*p* < 0.05). Additionally, RFP pixel density on the implant side was significantly higher during days 2-70 post-implantation (except day 56) compared to day 0 (*p* < 0.05). While contralateral tissue exhibited a small peak in RFP pixel density within the first week, no significant differences were detected between time points. This small peak transient in contralateral RFP pixel density indicates that the injury of the craniotomy window was minimal. Furthermore, the density of RFP clusters had a similar pattern to RFP pixel density over time, showing significant elevation during days 2-70 post-implantation on the implant sides versus the contralateral sides (*p* < 0.05, **Supplementary Fig. 2A-B**). Thus, the persistently elevated RFP activity near the microelectrode suggested an overall elevated autophagy activity throughout the 12-week implantation period, including a peak during the acute neuroinflammation period following a persistent elevation during the chronic foreign body response period.

Next, given the overall increase of autophagy vesicles, we examined if there was an increase in the immature autolysosome population. The presence of EGFP indicates the population of immature autophagy vesicles without acidification (autophagosomes). To examine changes in the population of immature autophagy vesicles near the microelectrode over time, the pixel density of EGFP was quantified at the implant and contralateral sides for each time point (**Fig. 2C**). Peaking at day 3, the EGFP pixel density on the implant side was significantly higher than contralateral side between days 2-17 post-implantation (*p* < 0.05). Of note, EGFP pixel density on the implant side became significantly elevated compared to the contralateral side from chronic days 63-77 post-implantation (*p* < 0.05). EGFP pixel density on the implant side was significantly elevated over acute (days 2-21 post-implantation) and chronic (days 70-77 post implantation) implantation periods compared to day 0 (*p* < 0.05). Moreover, the patterns of EGFP cluster density further confirmed the chronic accumulation of EGPF+ activity at the electrode-tissue interfaces (*p* < 0.05, **Supplementary Fig. 2C-D**). The contralateral EGFP pixel density showed a similar transient peak within the first week (*p* > 0.05), just as the contralateral RFP pixel density, which indicates minimal brain injury due to the craniotomy without an implant. Together, these results demonstrate a peak of autophagosome activity during the acute neuroinflammation period and a chronic buildup of immature autophagy population near the implanted microelectrode, suggesting a dysfunction of intracellular autophagy maturation near the microelectrode-tissue interface.

Given that there was a chronic accumulation of immature autophagosome population near the implant, we next investigated if these immature autophagosomes are able to acidify. To investigate the activity of acidified autolysosomes near the chronically implanted microelectrode, we looked at RFP+ EGFP fluorescent signals. Mature, acidified autophagy vesicles (autolysosomes) quench EGFP signals but retain RFP fluorescence. Thus, the pixel density of RFP+ EGFP signals were measured to quantify the population of mature autophagy vesicles at the implant and contralateral sides over time (**Fig. 2D**). We found the RFP+ EGFP pixel density was significantly higher on the implant side compared to the contralateral side for days 2-77 post-implantation (*p* < 0.05). Additionally, the RFP+ EGFP pixel density on the implant side was significantly elevated at days 2-77 post-implantation compared to day 0 (*p* < 0.05). Thus, a significant population of mature, acidified autophagy vesicles appeared throughout the long-term microelectrode implantation period. These findings suggest that implantation injury is accompanied by a persistent and robust elevation of acidified autolysosomes in local brain tissue near the microelectrode.

Given that we observed an increase in both immature autophagosomes and acidic autolysosomes, we next investigated the distribution of immature and mature autolysosomes. To investigate whether the autophagy load near the implanted microelectrode was dominated by the population of immature autophagy vesicles (autophagosomes), the ratio of EGFP pixel density over RFP pixel density was calculated over time (**Fig. 2E**). A greater ratio indicates that the majority of RFP+ autophagy population are EGFP+ immature autophagy vesicles. We found that this ratio value was significantly elevated on the implant side compared to the contralateral side at days 1-7 post-implantation (*p* < 0.05), suggesting that implantation injury led to a predominant population of immature autophagosomes during the acute neuroinflammation period. Additionally, the ratio value on the implant side was significantly higher on days 1 – 4 compared to day 0 (*p* < 0.05), implying that the predominant autophagosome population occurred within the first week of implantation. There was no significant difference in the ratio value on the contralateral side between time points. Thus, the autophagy activity near the acute implanted microelectrode was dominated by immature autophagy vesicles. While autophagy activity peaked during the acute neuroinflammation period (within 7 days post-implantation), there was an abnormal accumulation of immature autophagy vesicles near the chronically implanted microelectrodes, suggesting that the autophagy lysosomal pathway was impaired over the chronic implantation period.

Given our observation of distinct patterns of autophagy populations at acute neuroinflammation and chronic foreign body response period near the microelectrode, we next investigated morphological changes in the size of autophagy vesicle clusters. The size of autophagic clusters likely reflect the extent of autophagosome accumulation inside the cells. We observed that many clusters had glial-like appearances, especially those proximal to the implanted microelectrode during the acute neuroinflammation period (**Fig. 2F**), suggesting glial cells experienced elevated autophagy activity. The representative yellow/green cluster on day 3 post-implantation showed morphologies consisting of a large bright body and hazy-like processes (white arrows in insert). However, the morphologies of autophagy clusters shifted to small dot-like structures over the chronic foreign body response period (**Fig. 2G**), suggesting a gradual reduction in autophagy activity. The changes in cluster size matched the visual observation of autophagy cluster morphologies. The RFP clusters on the implant side were significantly larger than clusters on the contralateral side at days 4 – 9, 13, 17 post-implantation (*p* < 0.05, **Fig. 2H**). Additionally, the RFP clusters at electrode-tissue interfaces significantly increased in size at days 4-7 post implantation relative to day 0 (*p* < 0.05). EGFP cluster sizes were significantly larger at days 3-7 post-implantation for the implant side compared to time-matched contralateral side and to day 0 of the ipsilateral implant side (*p* < 0.05, **Fig. 2I**). Interestingly, the peak size of EGFP clusters (138.82 ± 85.02 μm^2^) at day 4 was much larger than the peak size of RFP clusters (67.47 ± 33.55 μm^2^) at day 7 post-implantation, which supports the finding that autophagosome populations dominates the acute neuroinflammation period. However, the sizes of EGFP and RFP clusters during chronic implantation was similar (RFP cluster: 36.44 ± 11.44 μm^2^, EGFP cluster: 34.56 ± 14.15 μm^2^, at 84-days post implantation). In summary, these findings suggest that the population and distribution of autophagy vesicles within cells are heightened in large-size, glial-like appearances during acute neuroinflammation period and then decreased in small, sphere-like particles over the chronic implantation period.

### 3.2. Autophagy activity proximal to the implanted microelectrode over time

During the acute neuroinflammation period, activated microglia quickly cover the implant surfaces and reactive astrocytes become hypertrophic, resulting in gliosis that encapsulates the implanted microelectrode. Evidence shows that autophagy modulates microglia inflammation and astrocyte reactivity [17, 18, 69, 70]. Thus, we first evaluated the changes in autophagy activity at the electrode-tissue interface over the implantation period. This activity was measured by quantifying RFP and EGFP within a tissue volume of 20 μm above the implant surface. The autophagy activity at the surface of the implant appeared to increase during the acute implantation period (**Supplementary Fig. 3A**) and gradually decrease over time (**Supplementary Fig. 3B**). Both RFP and EGFP density over the implant surface was significantly elevated at days 3-6 post-implantation compared to day 0 (*p* < 0.05, **Supplementary Fig. 3C-D**), which suggested that there was a transient elevation of immature autophagosomes population proximal to the implant occurs within the first week of implantation. Next, we investigated the acidified, mature autolysosomal activity proximal to the implant by examining the RFP+ EGFP density. The RFP+ EGFP density over the implant surface showed significant elevations at days 4–6, 9–10, 12, and 21 post-implantation compared to day 0 (*p* < 0.05, **Supplementary Fig. 3E**). Given there was a peak in autophagy activity on the implant surface during the acute inflammatory phase on day 3, we then asked whether this peak was dominated by the population of immature autophagosomes by looking at the ratio of EGFP density over RFP density. This ratio showed significant elevation at 3 days post-implantation compared to day 0 (*p* < 0.05, **Supplementary Fig. 3F**), indicating that autophagosome population is upregulated near the implant during the acute neuroinflammation period. Next, we examined the morphologies of autophagy clusters over the implant surface. The size of RFP and EGFP cluster over the implant surface peaked acutely followed by a decreasing trend over the chronic period (*p* > 0.05, **Supplementary Fig. 3G-H**). In summary, the density and morphologies of autophagy vesicles proximal to the implanted microelectrode demonstrated a transient peak during the acute neuroinflammation period and reduced to lower levels during chronic implantation period.

Given the temporal changes in autophagy activity in response to microelectrode implantation, we next examined the spatial distribution of autophagy activity near the microelectrode. To investigate the spatial distribution of autophagy activity near the implanted microelectrode over time, heatmaps of RFP and EGFP cluster were examined as a function of distance from the electrode over 84-days post-implantation. First, we examined the total autophagy population by examining the spatial distribution of RFP clusters. The spatiotemporal heatmap of RFP cluster counts demonstrated a transient elevation up to 300 μm away from the probe within the first 7-days post-implantation and an accumulation within 50 μm at days 21-70 post-implantation (**Fig. 3A**). Next, we examined the EGFP clusters as an indicator of immature autophagosomes population. Unlike RFP clusters, EGFP cluster heatmap had a transient elevation mainly at 0-50 μm distance from the implant within the first 7-days post-implantation (**Fig. 3B**). Then, the spatial distribution of autophagy clusters were investigated by measuring the changes in cluster counts at each spatial bin over implantation time. Within 0-50 μm distance from the implant, EGFP cluster count was significantly increased between days 3 – 5 post-implantation and RFP cluster count was significantly elevated and peaked at day 5 post-implantation compared to day 0 (*p* < 0.05, **Fig. 3B**). These findings further support the idea that total population of autophagy vesicles near the implant transiently peaks during the acute neuroinflammation period. Although no significant difference in cluster count between time points was detected at further distance, RFP cluster count generally peaked within the first 7-days post-implantation up to 300 μm away from the probe (*p* > 0.05, **Fig. 3D-H**), suggesting that implantation injury leaded to a transient increase of the total autophagy population at the interface as well as distal cortical tissue during acute neuroinflammation period. In contrast, few EGFP clusters were detected at distances beyond 50 μm, implying that the immature autophagosomes were mainly localized to the interface tissue. Together, these results demonstrated that autophagy activity primarily occurred proximal to the implanted microelectrode.

**Figure 3.**
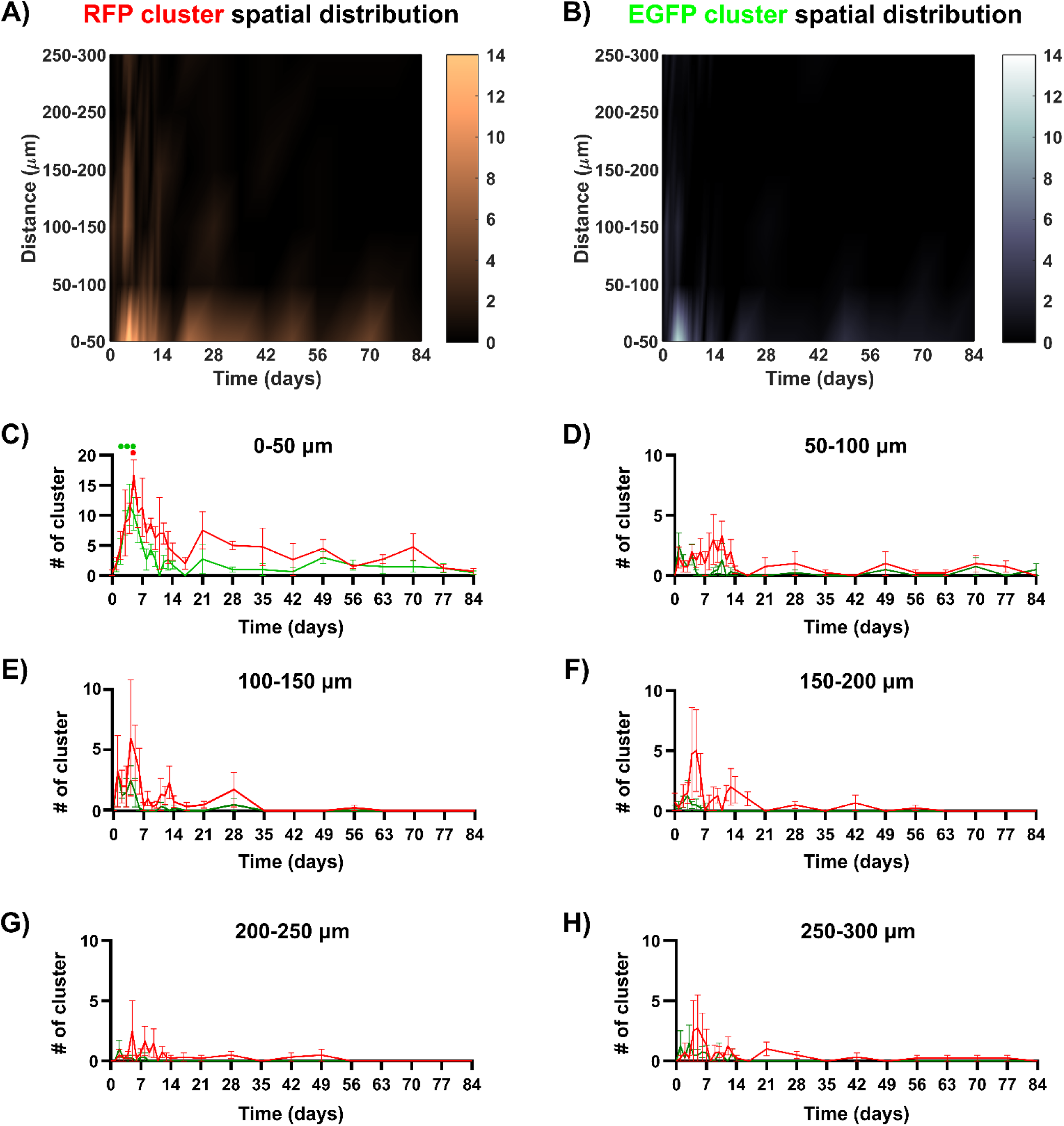
Proximal accumulation of autophagy activity near the microelectrode during chronic implantation. A spatiotemporal heatmap illustrates the number of RFP cluster **(A)** and EGFP cluster **(B)** at varying distances from the probe over the course of implantation. Visual observation revealed an increased number of RFP clusters and EGFP clusters in close proximity to the implant during the acute phase (days 0-14 post-implantation). **C)** The numbers of RFP clusters (red line) and EGFP clusters (green line) were plotted at 0-50 μm from the probe over microelectrode implantation period. The number of RFP cluster within 50 μm from the probe at day 5 post-implantation was significantly higher than the day 0. Similarly, the number of EGFP cluster within 50 μm at days 3-5 post-implantation was significantly higher than day 0. Few clusters of RFP and EGFP signals are observed in spatial bins of 50-100 μm **(D)**, 100-150 **(E)**, 150-200 **(F)**, 200-250 **(G)**, 250-300 **(H)**. Solid circles indicate significant differences in cluster numbers at the implant side between day 0 and other time points (One-way ANOVA followed by Dunnett’s test, *p* < 0.05). Red solid circle represents RFP clusters, while green solid circles represent EGFP cluster. All data were presented as mean ± standard deviation.

### 3.3. Distinct patterns of autophagy activity in different brain cells over long-term microelectrode implantation

Since two-photon imaging of CAG-LC3b-RFP-EGFP transgenic mice measured real-time autophagy activity independent of cell types, we used immunohistology to investigate changes in autophagy activity in different brain cells near the implanted microelectrode between 7- and 84-days post-implantation. The normalized intensities of LC3B autophagy markers were visibly elevated up to 300 μm away from the implant at 7-days post-implantation relative to 84-days post implantation (*p* > 0.05, **Fig. 4A**). The normalized intensities proximal to the implant was high at 84-days post-implantation relative to the distal tissue > 300 μm away from the implant. These immunohistology observations supported the results from two-photon imaging that implantation injury resulted in increased autophagy activity at interfaces and distal cortical tissue during the acute neuroinflammatory period.

**Figure 4.**
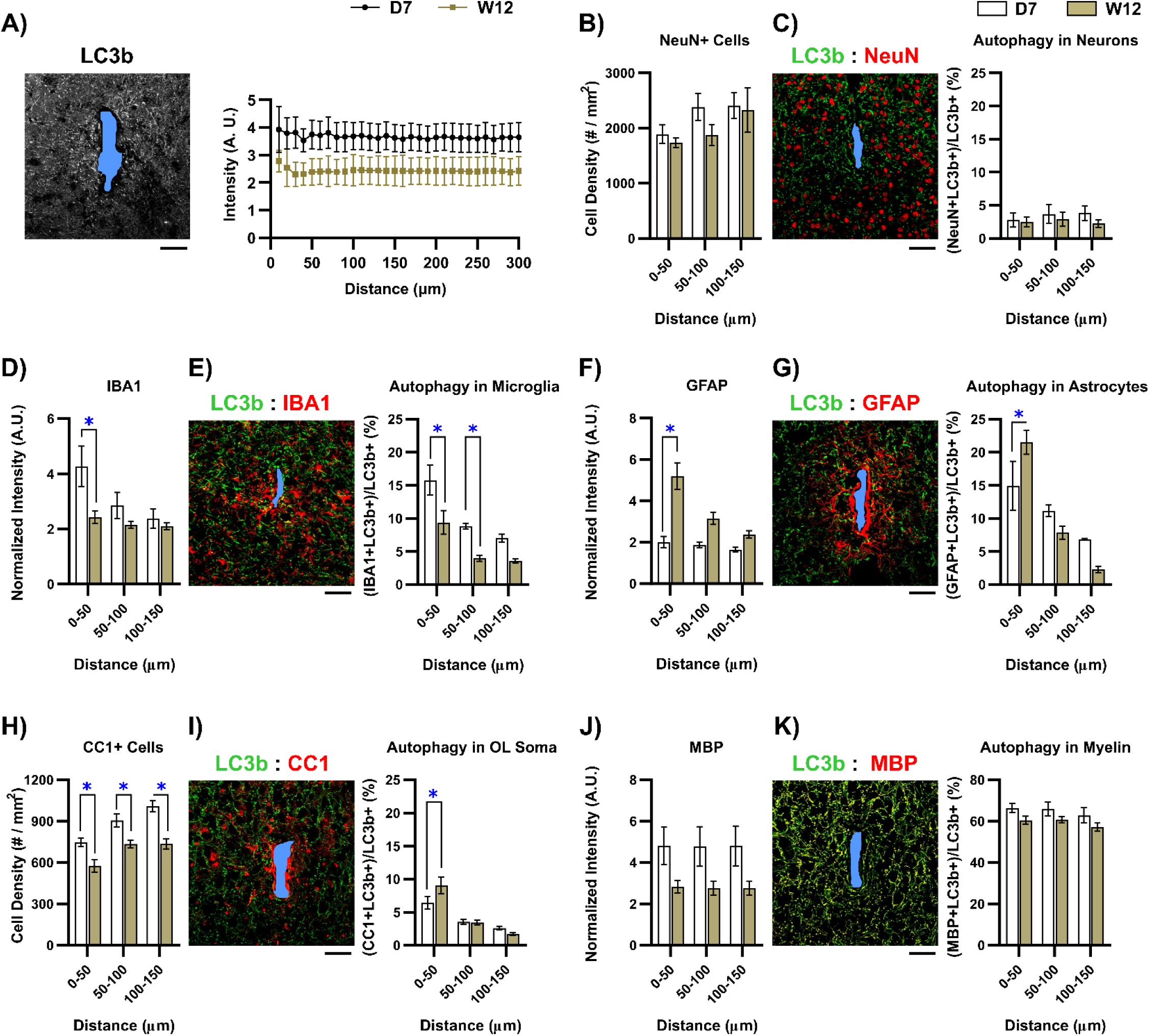
LC3b+ autophagy activity decreases in microglia, but increases in astrocyte and oligodendrocyte soma over time. **A)** Representative immunohistological staining of LC3b+ autophagy activity around the implant site on day 7 post-implantation. Intensity plot shows a global increase in LC3b+ autophagy activity on day 7 relative to day 84 post implantation. **B)** Density of NeuN+ cells showed a decreasing trend at 50-100 μm away from the electrode on day 84 post-implantation. **C)** The percentage of LC3b fluorescent signal colocalized with NeuN+ neurons were comparably low on day 7 and day 84 post-implantation at each spatial bin. **D)** Normalized intensity of IBA1 significantly decreased within 50 μm of the electrode on day 84 compared to day 7 post-implantation. **E)** Significant decrease in the percent colocalization of LC3b fluorescent signal with IBA1+ marker between day 7 and day 84 post-implantation at 0 – 100 μm distance bins indicates decreased microglia autophagy activity during chronic implantation. **F)** Normalized intensity of GFAP significantly increased near the implant on day 84 post-implantation. **G)** Significant increase in LC3b fluorescent signal colocalization with GFAP at 84-days post-implantation suggests enhanced autophagy activity in astrocyte near the probe during chronic implantation. **H)** CC1+ cell density significantly decreased between 0-150 μm from the implant on day 84 post-implantation. **I)** The percentage of LC3b fluorescent signal colocalized with the CC1+ oligodendrocyte soma significantly increased on day 84 compared to day 7 post-implantation at 0-50 μm. **J)** Normalized intensity of MBP showed a tendency to decrease near the implant on day 84 relative to day 7 post-implantation. **K)** No significance differences were detected in the high percentage of colocalizated LC3b and MBP+ myelin between day 7 and day 84 post implantation. Scale bar = 50 μm. Blue asterisks indicate significant differences in colocalization percentage between time points at each distance bin (Two-way ANOVA with Tukey post-hoc, *p* < 0.05). All data were presented as mean ± standard deviation.

Given the accumulation of immature autophagy vesicles during the chronic foreign body response phase, we then asked which cells increased their intracellular autophagy activity near the implanted microelectrode over time. The percentage of LC3B autophagy markers colocalized with different cellular markers (NeuN/IBA1/GFAP/CC1/MBP) was quantified at the acute 7-day and chronic 84-day post-implantation. We first observed that NeuN+ cell density experienced a decreasing trend at 50-100 μm away from the implant on day 84 post-implantation (**Fig. 4B**), suggesting a progressive degeneration of neuronal soma at chronic foreign body response phase. Neuronal autophagy, LC3B+ NeuN+ pixels, accounted for ∼5- 9% of all LC3B+ pixels and did not significantly change over time (*p* > 0.05, **Fig. 4C**).

Next, we examined cell types involved in glial scarring. Normalized intensity of microglia marker IBA1 was elevated proximal to the implant on day 7, then showed a significant decrease at 0-50 μm away from the implant by day 84 post implantation (*p* < 0.05, **Fig. 4D**). The intensity of autophagy LC3B markers within IBA1+ microglia at 0-50 μm distance from the implant significantly decreased from 16% at day 7 to 9% at day 84 post-implantation (*p* < 0.05, **Fig. 4E**). In contrast, GFAP normalized intensity significantly increased 0-50 μm away from the implant on day 84 compared to day 7 post-implantation (*p* < 0.05, **Fig. 4F**). The percentage of autophagy LC3B markers within GFAP+ astrocytes at 0-50 μm distance from the implant significantly increased from ∼15% on day 7 post-implantation to ∼21% on day 84 post implantation (*p* < 0.05, **Fig. 4G**).

Lastly, we investigated autophagy activity in oligodendrocyte lineage structures. Although the population of CC1+ oligodendrocyte soma significantly decreased at the implant interface (*p* < 0.05, **Fig. 4H**), the percentage of autophagy LC3B markers within CC1+ oligodendrocyte soma at 0-50 μm distance from the implant significantly increased at 84-days post-implantation (∼ 6% at 7-days, ∼ 9% at 84-days, *p* < 0.05, **Fig. 4I**). The normalized intensity of MBP staining showed a decreasing trend of myelin density near the implanted microelectrode at 84-days post-implantation (**Fig. 4J**), suggesting a progressive demyelination at chronic foreign body response phase. Despite that no significant difference was detected between time points at each spatial bin, the percentages of autophagy LC3B markers with MBP+ myelin near the implanted electrode were highly visible (∼55-63% of all LC3B+ pixels) (**Fig. 4K**). Together, we observed different patterns of changes in autophagy vesicle population within different cell types over the long-term microelectrode implantation. Specifically, microglial autophagy decreased but astrocytic and oligodendrocyte soma autophagy increased proximal to the chronically implanted microelectrode.

### 3.4. Impairment in neuronal lysosome activity near the chronically implanted microelectrode

Lysosome fusion with autophagosomes is a critical process for the autophagy degradation [71]. Given the observation that autophagy was disrupted over the chronic implantation period, we next asked if lysosomes were impaired around the implanted microelectrode. To investigate lysosome activity, we characterized the changes in autophagy-lysosomal degradation by staining for LAMP1, a lysosomal marker, at days 7, 14, 28, and 84 post-implantation (**Fig. 5A**). Normalized LAMP1 intensities appeared to be higher proximal to the implant side at days 14 and 84 post-implantation (**Fig. 5B**). The average normalized LAMP1 intensity significantly decreased within 50 μm distance from the implant on day 28 and significantly elevated by day 84 post-implantation (*p* < 0.05, **Fig. 5C**). These results suggested that lysosome activity following the implantation were acutely elevated during the inflammatory phase and then accumulated around the implanted microelectrode over the chronic foreign body response phases.

**Figure 5.**
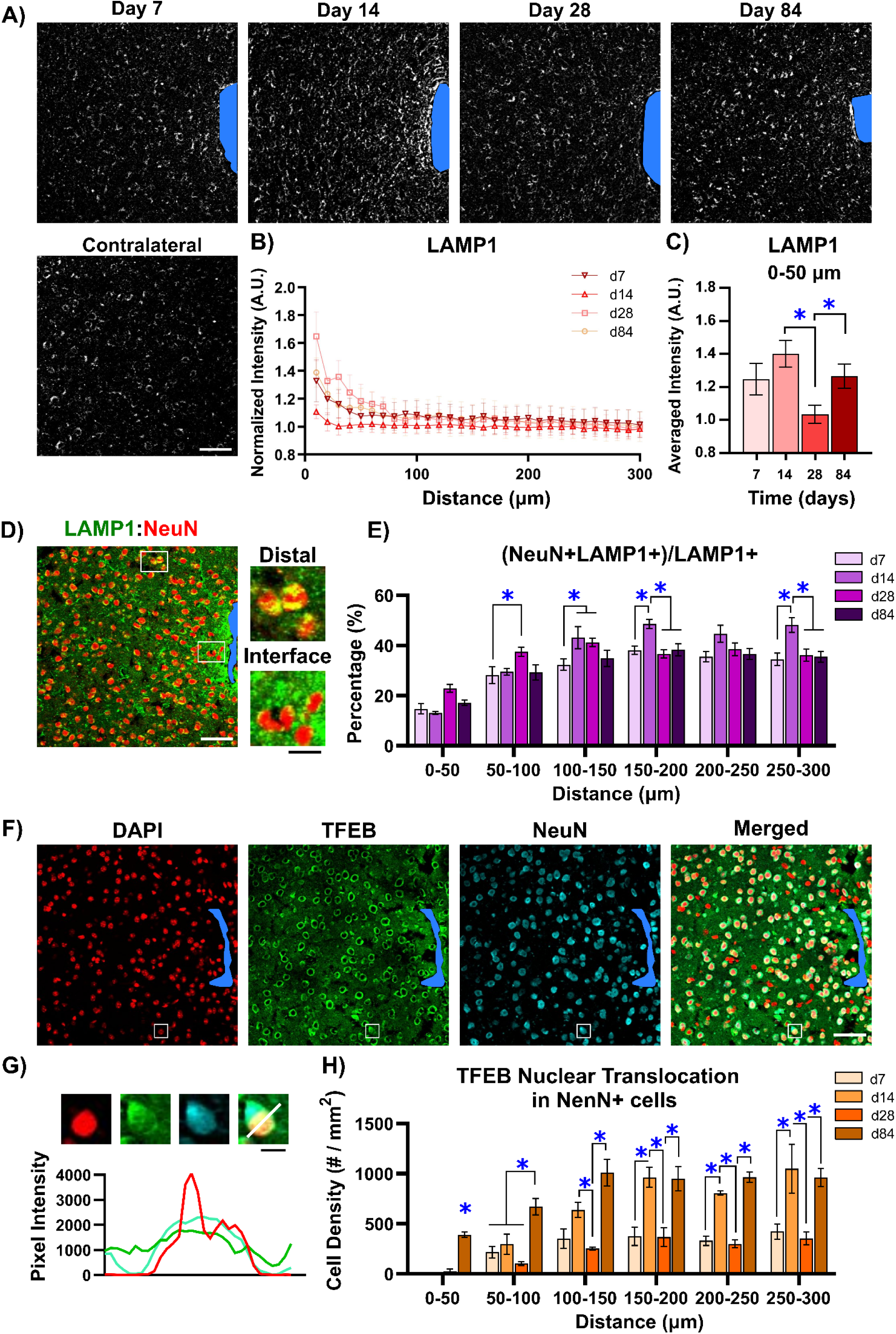
Abnormal lysosomal activity in neurons near the chronically implanted microelectrode. **A)** Representative images showing LAMP1+ lysosome staining around the implanted microelectrode at days 7, 14, 28, and 84 post-implantation, as well as at the no-implant contralateral side on day 84 post-implantation. Scale bar = 50 μm. **B)** Normalized LAMP1 fluorescent intensity plotted as a function of distance near the implanted microelectrode on days 7, 14, 28, and 84 post-implantation. **C)** Averaged LAMP1 fluorescence intensities significantly decreased on day 28 and then significantly increased on day 84 post-implantation. **D)** Immunohistological representation showing colocalization of LAMP1+ lysosome vesicles with NeuN+ neuronal soma around the microelectrode implant site on day 7 post implantation. Magnified insert visually demonstrate that neurons at interface have fewer lysosome vesicles compare to neurons at distal regions. White scale bar = 50 μm. Black scale bar = 10 μm. **E)** Percentage of LAMP1 fluorescent signals colocalized with NeuN+ was measured within 50 μm bins up to 300 μm from the implant site on days 7, 14, 28, and 84 post-implantation. **F)** Representative image showing immunohistological staining of nuclei (DAPI), transcription factor EB (TFEB), neuronal soma (NeuN) on day 84 post-implantation. **G)** TFEB nuclear translocation in neurons detected by triple fluorescence overlap in DAPI (red), TFEB (green), and NeuN (cyan) channels, suggesting lysosomal biogenesis in neurons. Scar bar = 10 μm. **H)** Density of TFEB nuclear translocation cells colocalized with NeuN markers within 50 μm bins up to 300 μm from the implant site. Blue asterisks indicate significant differences in quantification metrics between different time points at each distance bin (Two-way ANOVA with Tukey post-hoc, *p* < 0.05). All data were presented as mean ± standard deviation.

Having observed the changes in the lysosomal population in cortical tissue near the implant, we next investigated the cell specific changes in lysosome activity. Neuronal loss has been observed near the chronically implanted microelectrode on day 84 post-implantation, indicating an increased neuronal degeneration within 100 μm away from the electrode (**Supplementary Fig. 4A**). To determine changes in neuronal lysosome load around the microelectrode over time, the percentages of LAMP1 marker colocalized with NeuN+ neurons were measured in 50 μm bins up to 300 μm distance from the implant at days 7, 14, 28, and 84 post-implantation. We observed that LAMP1 colocalization with NeuN+ cells decreased at the close electrode-tissue interface (**Fig. 5D**). However, LAMP1 colocalization with NeuN+ neurons significantly increased at 100-300 μm distance from the implant site on day 14 post-implantation (*p* < 0.05, **Fig. 5E**). The reduced lysosome load in neurons proximal to the implant (**Fig. 5E**, **Supplementary Fig. 4B**) inspired us to investigate whether the microelectrode implantation injury disrupts lysosomal biogenesis. We co-stained DAPI, Transcription factor EB (TFEB), NeuN and measured the density of neurons with TFEB nuclear translocation (**Fig. 5F**). TFEB is a master regulator of autophagy-lysosomal pathway, and TFEB dephosphorylation and nuclear translocation promote the expressions of genes codding for lysosomal proteins [72, 73]. Neurons with TFEB nuclear translocation were identified by the triple fluorescent colocalization of DAPI, TFEB, and NeuN (**Fig. 5G**; for neurons without TFEB translocation see **Supplementary Fig. 4C**). We observed the densities of neurons with TFEB nuclear translocation were significantly elevated in tissue 150-300 μm away from the implant site on day 14 post-implantation. Interestingly, we also found that the densities of neurons with TFEB nuclear translocation were significantly elevated up to 300 μm distance from the implant at chronic day 84 compared to day 28 post-implantation (*p* < 0.05, **Fig. 5H**), suggesting increased lysosomal biogenesis at distal tissue over the chronic implantation period. However, the percentage of LAMP1 markers within NeuN+ neurons was comparable between days 28 and 84 post-implantation (**Fig. 5E**), indicating that the lysosomal population did not increase during the chronic implantation period. Together, these findings revealed that chronic implantation resulted in a mismatch between the lysosome population and lysosomal biogenesis in neurons near the microelectrodes.

### 3.5. Elevated lysosome load and myelin debris clearances in reactive astrocytes proximal to the implanted microelectrode

While microelectrode implantation injury disrupts neuron viability, glial morphologic changes are also observed near the implanted microelectrode [17, 18, 30]. Lysosomes play a critical role in regulating glial activity and morphological changes under pathological conditions [74, 75]. To examine the relationship between lysosome activity and glial cells near the microelectrode, we measured the percentage of LAMP1 markers colocalized with glial markers (IBA1/GFAP/CC1/MBP) in 50 μm bins up to 300 μm distance from the implant site on days 7, 14, 28, and 84 post-implantation.

Microglia and astrocytes are involved in glial scarring during acute neuroinflammation and become relatively morphologically stable during chronic foreign body response [14]. IBA1+ microglia intensities demonstrated significant elevations within 0-100 μm away from the microelectrode at acute 0-14 days post implantation (**Fig. 6A**; *p* < 0.05, **Fig. 6B**). The percentages of LAMP1 vesicles colocalized with the IBA1+ microglia significantly dropped 0-200 μm tissue away from the implant site starting on 28-days post-implantation (*p* < 0.05, **Fig. 6C**), which suggested the microglia has an elevated lysosome load proximal to the implant during acute neuroinflammatory period compared to chronic foreign body response phase.

**Figure 6.**
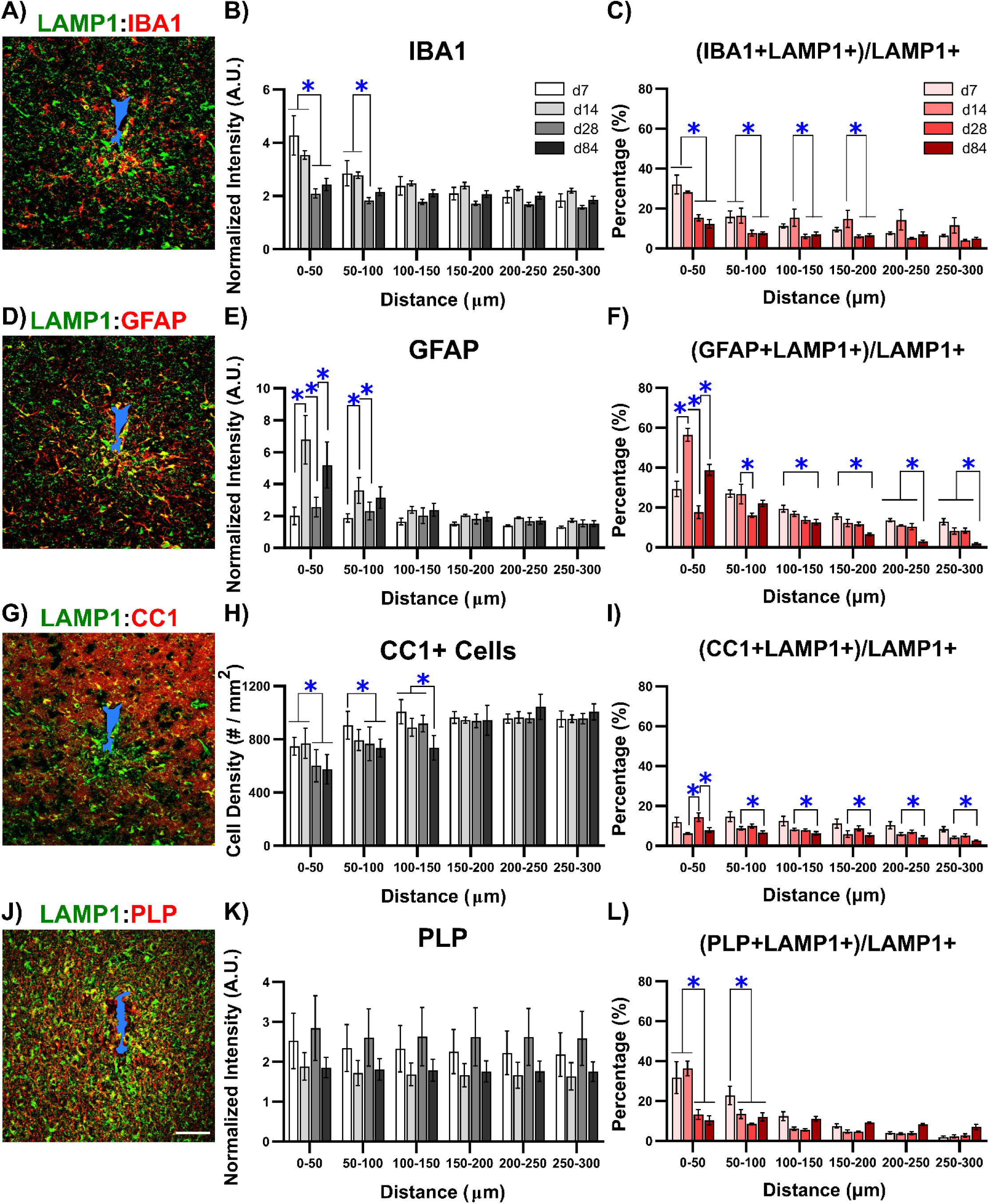
Increased lysosomal activity in astrocytes near the microelectrode during chronic implantation. **A)** Colocalization of LAMP1 and IBA1 staining. **B)** Significant elevations of IBA1 intensity occurred within 100 μm from the microelectrode on days 7 and 14 post-implantation. **C)** A significant decrease in the percentage of LAMP1 fluorescent signal colocalized with IBA1+ microglia was observed on day 28 post-implantation within 0-100 μm from the implant, indicating increased lysosome activity in microglia near the implant during acute implantation period. **D)** Colocalization of LAMP1 and GFAP markers. **E)** GFAP+ intensity peaked on day 14 and increased on day 84 post-implantation within 50 μm of the implant. **F)** The percentage of LAMP1 signal colocalized with GFAP+ reactive astrocytes within 50 μm significantly increased on day 14, then decreased by day 28, and eventually increased again on day 84 post-implantation. **G)** Colocalization of LAMP1 and CC1 markers near the implanted microelectrode. **H)** CC1+ cell density significantly decreased on day 28 within 100 μm of the implant and on day 84 between 100-150 μm from the implant. **I)** The percentage of LAMP1 fluorescent signals colocalized with CC1+ oligodendrocyte soma between 50-300 μm from the implant showed significantly low level of lysosome activity on day 84 compared to day 7 post-implantation. **J)** Colocalization of LAMP1 and PLP marker near the implant. **K)** PLP intensity tended to decrease on day 84 post implantation. **L)** The percentage of LAMP1 signals colocalized with PLP+ myelin showed a significantly higher percentage within 50 μm of the probe on day 7 and day 14 post-implantation compared to day 28 and day 84 post-implantation. White scale bar = 50 μm. Blue asterisks indicate significant differences in colocalization percentage between 7- and 84-days post implantation at each distance bin (Two-way ANOVA with Tukey post-hoc, *p* < 0.05). Data are presented as mean ± standard deviation.

GFAP+ astrocyte activity significantly peaked on day 14 post-implantation (**Fig. 6D**; *p* < 0.05, **Fig. 6E**). Additionally, the percentages of LAMP1 markers colocalized with the GFAP+ astrocytes showed a significant, but transient peak (56.49 ± 4.53 %) at 0-50 μm distance from the implant on day 14 post-implantation. Interestingly, the percentages of astrocyte LAMP1 markers colocalized with GFAP+ significantly elevated again (38.83 ± 6.21 %) at 0-50 μm away from the implant on day 84 post-implantation. Yet, the percentages of LAMP1+GFAP+ pixels at 100-300 μm tissue away from the implant site gradually decreased over time (*p* < 0.05, **Fig. 6F**). These findings suggested that 1) lysosomal activity in microglia and astrocytes experienced a transient increase within the first two weeks of implantation corresponding to the acute neuroinflammation phase (*p* < 0.05, **Supplementary Fig. 5A-B**); 2) astrocytic lysosomes was chronically elevated proximal to the implanted microelectrode possibly contributing to the chronic accumulation of the lysosome populations over the chronic implantation period.

Having observed increased lysosomes load in microglia and astrocytes for glia scars, we next asked whether lysosomal activity in oligodendrocytes and myelin were influenced by long-term implantation. Lysosomes regulate myelin integrity, sheath growth, and oligodendrocyte differentiation [76–78]. However, implantation injury leads to progressive oligodendrocyte loss and demyelination within 50 μm away from the implant (**Fig. 6G**; *p* < 0.05, **Fig. 6H**) [27]. Therefore, we asked if lysosomes also increased in the oligodendrocyte and myelin injury near the chronically implanted microelectrodes. The percentages of LAMP1 markers colocalized with the CC1+ oligodendrocytes showed a small significant peak at 0-50 μm away from the implant on day 28 post-implantation yet decreased between 50-300 μm away from the probe over time (*p* < 0.05, **Fig. 6I**). Additionally, the percentages of CC1 markers colocalized with the LAMP1 significantly decreased from 0-100 μm and 150-300 μm on day 84 post-implantation (*p* < 0.05, **Supplementary Fig. 5C**). These findings suggest lysosomal populations decreased in oligodendrocyte soma during the chronic implantation period.

The staining result for myelin marker PLP demonstrated a decreasing trend in PLP+ intensity on day 84 post-implantation (**Fig. 6J-K**). Of note, the percentages of LAMP1 markers colocalized with the PLP+ myelin were prominent within 0-50 μm away from the implant on day 7 (31.67 ± 11.31 %) and day 14 (36.27 ± 5.13 %) post-implantation. The LAMP1+PLP+ percentage significantly dropped within 0-50 μm from the implant beginning day 28 post-implantation and within 50-100 μm beginning day 14 post-implantation (*p* < 0.05, **Fig. 6L**; *p* < 0.05, **Supplementary Fig. 5D**). We found colocalization between PLP+ myelin cluster and GFAP+ LAMP1+ astrocytic lysosomes by examining the day 7 post-implanted brain tissue co-stained for LAMP1, PLP+, GFAP+, (**Fig. 7**, white arrows). These triple fluorescent colocalizations also suggest that astrocytes contributed to the myelin debris clearance via the autophagy-lysosomal pathway proximal to the electrode tissue interfaces during the acute implantation period. These results support that astrocyte lysosome activity was predominant near the microelectrode during the chronic implantation period.

**Figure 7.**
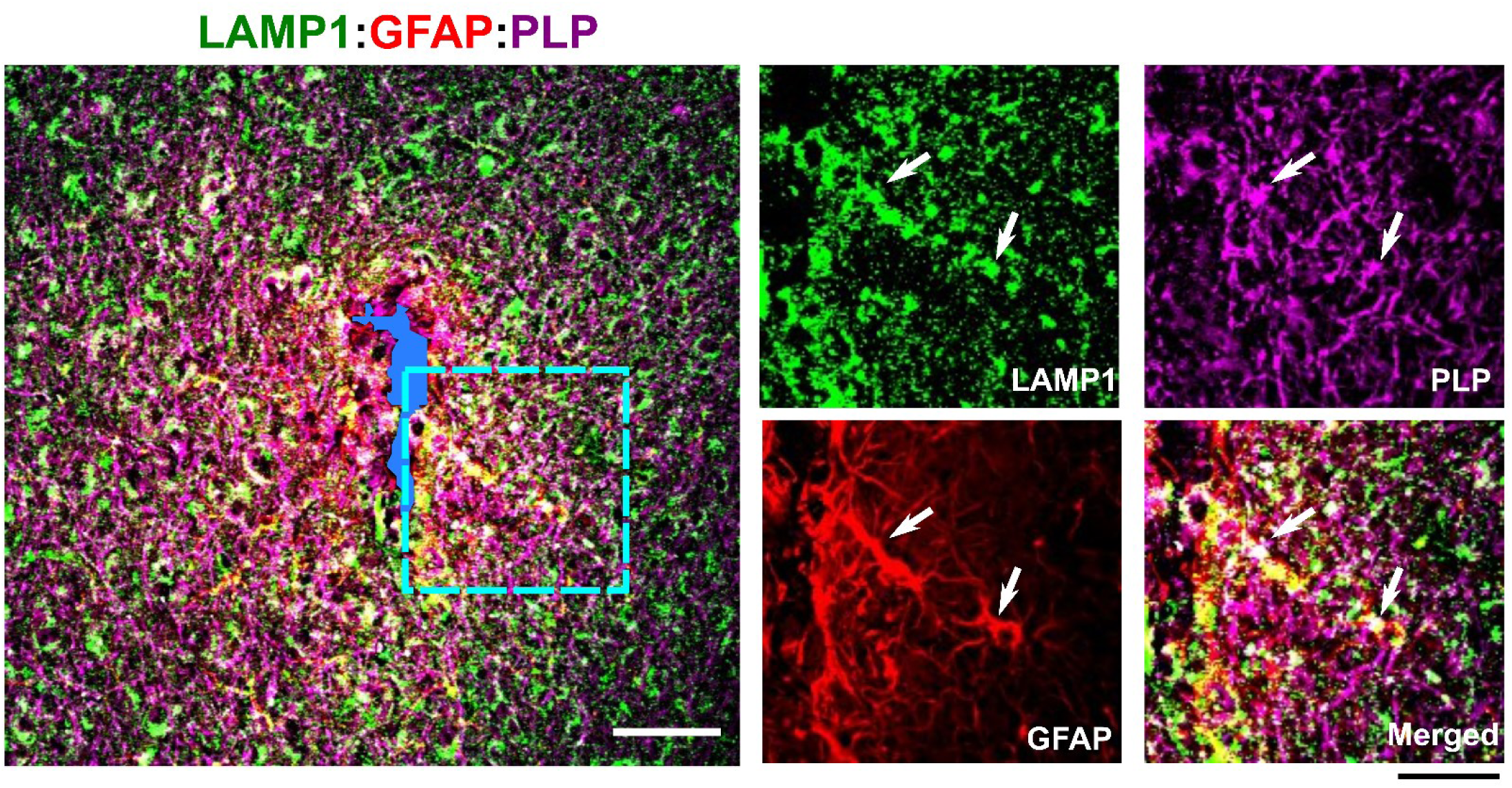
Astrocytic lysosomes involved in myelin debris clearance. Immunohistological representation of lysosomal vesicles (LAMP1, green) containing myelin protein (PLP, magenta) within reactive astrocytes (GFAP, red) near the implant site on day 7 post implantation, indicating the involvement of reactive astrocyte in degradation of myelin debris during acute implantation period. While scale bar = 50 μm. Black scale bar = 20 μm for image inserts.

### 3.6. Abnormal accumulation of autophagy cargo, decrease in a lysosomal protease, and accumulation of oxidative stress marker near the chronically implanted microelectrode

Given the abnormal autophagy-lysosomal activity near the chronically implanted microelectrode, we next investigated the associated pathologies. We first stained for an autophagy cargo marker SQSTM1/p62 and a lysosomal protease marker cathepsin D on brain tissue explanted on day 7, 14, 28, and 84 post-implantation. SQSTM1/p62 is a cargo receptor to recognize specific targets for autophagosome sequestration and eventual degradation by lysosomes. Thus, the accumulation of SQSTM1/p62 markers reflects autophagy blockade, which was visually elevated near the implant site on day 14 and day 84 post-implantation (**Fig. 8A**). The average normalized SQSTM1/p62 intensity significantly increased near the implant site (0-50 μm) on chronic day 84 post-implantation (*p* < 0.05, **Fig. 8B**), suggesting that autophagy lysosomal clearance was impaired near the chronically implanted microelectrode. Similarly, normalized cathepsin D intensities were elevated proximal to the implant site over the long-term implantation period (**Fig. 8C**). However, the average normalized cathepsin D intensities significantly decreased in tissue regions up to 150 μm away from the implant site from day 28 post-implantation (*p* < 0.05, **Fig. 8D**), suggesting deficits in lysosomal capability for autophagy cargo degradation near the chronically implanted microelectrode.

**Figure 8:**
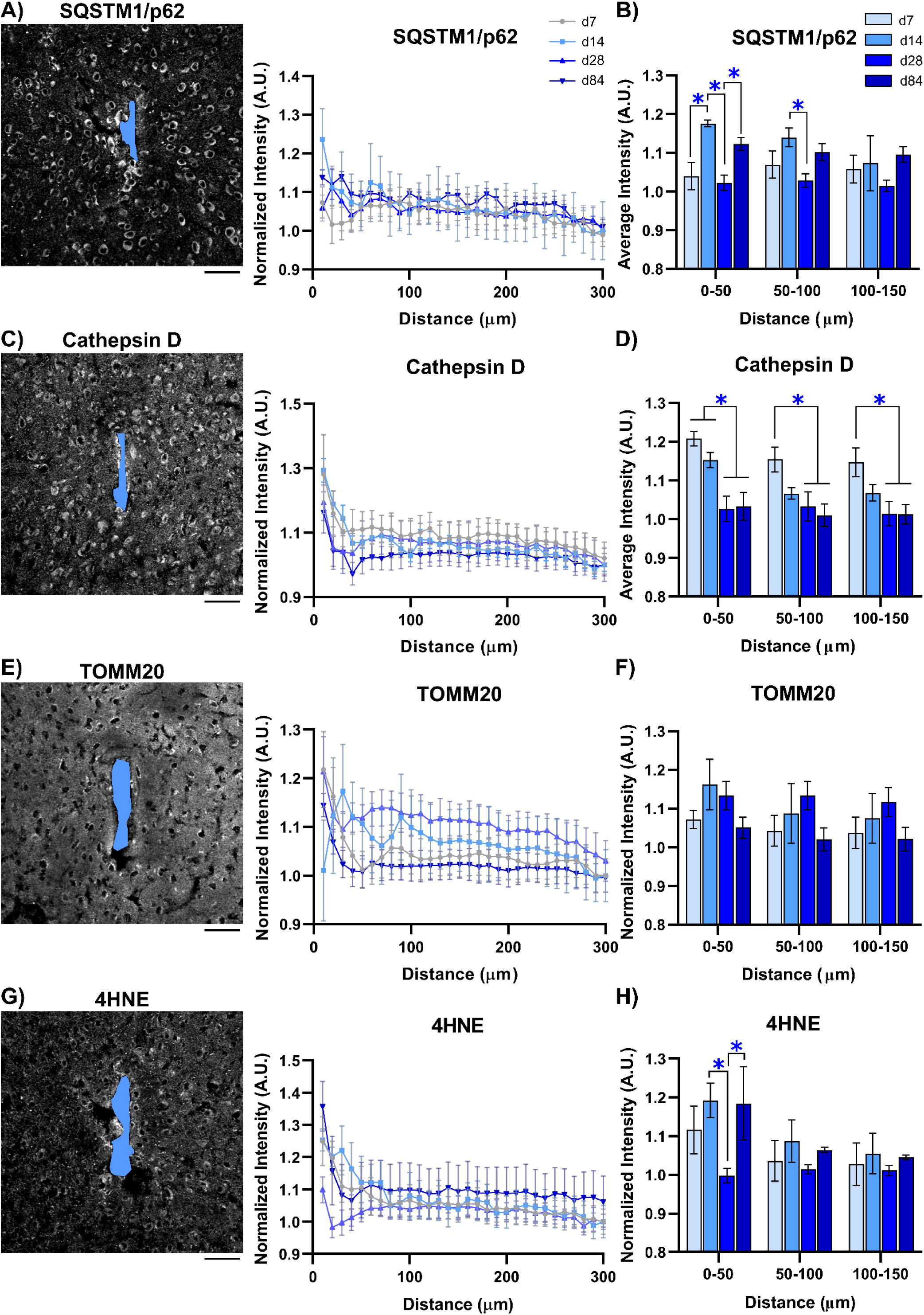
Accumulated autophagy cargo, reduced lysosomal protease, and increased oxidative stress at chronic implantation period. **A)** Normalized fluorescence intensity or autophagy cargo marker (SQSRM1/p62) plotted against distance from the implant site on days 7, 14, 28, and 84 post-implantation. **B)** A significant increase in averaged SQSTM1/p62 intensity was observed 0-50 μm from the electrode on day 84 post-implantation, suggesting the accumulation of autophagy cargo near the implant during the chronic implantation period. **C)** Normalized fluorescence intensity for acidic lysosomal protease (cathepsin D) plotted against distance from the implant site on days 7, 14, 28, and 84 post-implantation. **D)** Average cathepsin D fluorescence intensity was significantly lower within 150 μm of the probe on day 28 and 84 post-implantation compared to day 7, suggesting impaired lysosomal activity for intracellular degradation at the chronic implant interfaces. **E)** Normalized fluorescence intensity for mitochondria (TOMM20) plotted against distance from the implant site on days 7, 14, 28, and 84 post-implantation. **F)** Average TOMM20 fluorescence intensity within 50 μm bins up to 150 μm from the implant site on days 7, 14, 28, and 84 post-implantation. **G)** Normalized fluorescence intensity for oxidative stress (4-Hydroxynonenal (4-HNE)) was measured as a function of distance from the implant site on days 7, 14, 28, and 84 post-implantation. **H)** Average TOMM20 fluorescence intensity within 50 μm of the implant was significantly higher on day 84 post-implantation compared to day 28, indicating the accumulation of oxidative stress during the chronic implantation period. Scale bar = 50 μm. Blue asterisks indicate significant differences in colocalization percentage between 7- and 84-days post-implantation at each distance bin (Two-way ANOVA with Tukey post-hoc, *p* < 0.05). All data are presented as mean ± standard deviation.

Having observed deficits in the lysosomal protease that is required for autophagy near the chronically implanted microelectrode, we next investigated the mitochondria integrity and oxidative stress by staining for mitochondria marker (TOMM20) and 4-Hydroxynonenal (4-HNE), respectively. The profile of normalized TOMM20 intensities demonstrated visible decreases proximal to the electrode-tissue interfaces on day 84 post-implantation (**Fig. 8E-F**), suggesting the impairment of mitochondria activity around the chronically implanted microelectrode. Inefficient removal of damaged mitochondria leads to the accumulation of reactive oxygen species. Accordingly, the intensity of stain for oxidative stress marker 4-hydroxyninenal (4HNE) was elevated in the tissue proximal to the chronic implant(**Fig. 8G**). The average normalized 4HNE intensity showed a significant increase within 0-50 μm from the implant site on day 84 post-implantation (*p* < 0.05, **Fig. 8H**), suggesting a chronic accumulation of oxidative stress. Together, these results suggested that impaired clearance of autophagy cargo, insufficient lysosomal protease, and accumulation of oxidative stress contribute to the pathological autophagy-lysosomal degradation near the implanted microelectrode.

## 4. Discussion

The current challenge with brain microelectrodes in clinical applications includes the chronic decline in their functional performances [6, 8]. Tissue degeneration around the chronically implanted microelectrode is associated with this functional decline in stability and sensitivity of the recorded neural signals [79]. This neuronal loss, glial activation, vascular injury, and demyelination are also present in neurodegenerative diseases [14]. A growing number of studies reveal similar pathologies between the focal brain trauma by microelectrode implantation and neurodegenerative diseases [9, 27, 80, 81]. The intracellular autophagy-lysosomal pathway plays a vital role in the homeostasis and function of brain tissue, and deficits of this pathway have been observed in the onset and progression of neurodegenerative diseases such as AD [47, 49]. Therefore, we aimed to answer whether this autophagy-lysosomal pathway is impaired near microelectrodes that are implanted over time. Here, two-photon imaging showed an aberrant accumulation of immature autophagy vesicles near the chronically implanted microelectrode. Additionally, immunohistology revealed a discrepancy between elevated lysosomal population and the low protease capacity near the implant further indicating an aberrant accumulation of autophagy cargo. Our results help build a more comprehensive understanding of intracellular autophagy lysosome processes over the chronic microelectrode implantation period. These results highlight the understudied role of metabolic waste removal on tissue degeneration at neural interfaces, which may uncover new understanding and innovations for improving chronic tissue-electrode interfaces.

### 4.1. Impaired autophagy-lysosomal pathway is associated with the disease-associated factors following chronic microelectrode implantation

The autophagy-lysosomal pathway is a critical process for degrading harmful protein aggregates that appear to be cytotoxic [39, 82]. The abnormal accumulation of these protein aggregates, such as misfolded tau proteins and amyloid-β peptides, are characteristics of neurodegenerative diseases [83]. In this study, we show that chronically implanting microelectrodes results in an accumulation of immature autophagy vesicles (**Fig. 2C**), a local increase in SQSTM1/p62 labeled autophagy cargo (**Fig, 8A-B**), and a reduced level of lysosomal protease (**Fig. 8C-D**). These observations suggest that protein aggregates in autophagy vesicles are unable to achieve timely degradation near chronically implanted microelectrodes. In other words, the autophagy-lysosomal pathway is likely impaired throughout the chronic implantation period. Dysregulation and inefficient acidification of lysosomes may lead to the conversion of the autophagy cargo into lipofuscin and result in increased lipofuscin deposition. Moreover, the impaired clearance in pathological protein aggregates has been speculated as a critical factor of neurodegenerative diseases. For example, abnormal accumulation of SQSTM1/p62 markers and LC3B autophagy markers were colocalized with hyperphosphorylated tau in the brain tissue of Alzheimer’s disease patients [10], suggesting that the autophagy-lysosomal pathway plays a role in the clearance of pathological tau. Interestingly, a recent study has shown evidence of hyperphosphorylated tau near chronically implanted microelectrodes [9]. Therefore, it is possible that hyperphosphorylated tau is associated with the accumulation of impaired autophagy vesicles near chronically implanted microelectrodes. However, it remains to be seen whether the hyperphosphorylated tau is colocalized with SQSTM1/p62+ autophagy vesicles near the chronically implanted microelectrodes or whether the impaired autophagy-lysosomal pathway is a cause of hyperphosphorylated tau accumulation. Future work onunderstanding the role of autophagy-lysosomal degradation on tau pathologies would help to identify potential targets to mitigate the neurodegeneration due to the long-term implantation.

Besides protein aggregates, damaged mitochondria producing ROS are also degraded by the autophagy-lysosomal pathway in healthy tissue [84, 85]. ROS including free radicals, oxygen anions, and hydrogen peroxide, are by-products of the oxygen metabolism [85]. Increases in ROS serves as crucial signals to induce autophagy-lysosomal degradation to avoid oxidative injury [86–88]. In this study, we found a transient elevation of ROS near the implant within the first two weeks post-implantation (**Fig. 8G-H**), which corresponds to a peak autophagy activity at the electrode-tissue interfaces during the inflammatory period (**Fig. 2**). The normalized TOMM20 intensities didn’t not experience a significant drop on days 14 and 28 post-implantation (**Fig. 8E-F**), suggesting that mitochondria integrity is not severely damaged near the implanted microelectrode. These findings indicate that the peak autophagy activity may be a protective mechanism near the implant during acute implantation period. However, prolonged, excessive ROS accumulation results in oxidative stress, which can lead to irreversible damage to proteins, lipids, and even DNA during the pathogenesis of brain trauma and neurodegenerative diseases [44, 89–91]. Here, we demonstrate that 84-days microelectrode implantation can induce the oxidative stress near the implant (**Fig. 8G-H**). In turn, this oxidative stress is likely associated with the damaged population of mitochondria (**Fig. 8E-F**), which may not be efficiently removed via an impaired autophagy-lysosomal pathway. On the other hand, autophagy-lysosomal degradation is also vulnerable to oxidative stress [92–94], based on evidence that oxidative stress induces the permeabilization of the lysosomal membrane and thereby leads to neuronal death [95, 96]. Therefore, it is still unclear whether the oxidative stress is the consequence or cause of the impaired autophagy-lysosomal pathway around chronically implanted microelectrodes. Further evidence is needed to reveal the relationship between the oxidative stress and autophagy-lysosomal degradation to better understand the failure mechanisms of brain tissue at the neural interface.

### 4.2 Impaired microglial and astroglial autophagy-lysosomal pathway influences the phagocytosis following chronic microelectrode implantation

Microglia are key regulators of inflammatory responses around long-term implanted microelectrode as well as in the neurodegenerative diseases [18, 97]. During the neuroinflammatory phase, microglia become activated and contribute to debris clearances of damaged cells via the phagocytosis [98]. Phagocytosis is a mechanism for recognizing, engulfing, and degrading cells and extracellular materials in lysosomes [99]. Recently, an involvement of autophagy in this process has been demonstrated LC3-associated phagocytosis (LAP), in which autophagy machinery is partially translocated to the phagocytosis vesicles and results in intracellular processing of the engulfed extracellular cargo via lysosomal fusion [99–101]. Here, our findings demonstrate that microglia proximal to the implant increased in autophagy and lysosomal activity (**Fig. 4E**, **Fig. 6C**), indicating LAP-mediated clearance of implantation-damaged cells. However, the autophagy-lysosomal activity in microglia gradually decreases over time (**Fig. 4E**, **Fig. 6C**). Similar decline of microglial autophagy-lysosomal activity occurs in aging and neurodegenerative diseases such as AD, Parkinson’s Disease, which is associated with the impaired clearances of lipid droplets, amyloid β, and tau aggregates [20, 102–104]. With increased density of apoptotic neurons at chronic 16-weeks post-implantation [9, 33], the observation of reduced autophagy-lysosomal activity in microglia indicates an impaired phagocytic clearance. Yet, direct evidence is still lacking to confirm whether microglial phagocytic activity is impaired near the chronic microelectrode or which role autophagy plays in microglial phagocytosis. Future studies should clarify the relationship between impaired autophagy-lysosome activity within microglia and their phagocytic activity in both disease and brain trauma.

Similar to microglia, astrocytes become reactive and demonstrate a hypertrophic morphology following the microelectrode insertion [17, 105]. These reactive astrocytes together with microglia form an encapsulating glial scar around the implanted microelectrode over time [17]. Astrocytes, like microglia, also have the phagocytic capability to engulf of cellular debris, which was shown in models of brain injury and neurodegenerative diseases [106–108]. Reduced autophagy in astrocytes is associated with a compromised astrocytic clearance of amyloid in AD mice, suggesting the role of autophagy-lysosomal pathways in the astrocytic phagocytosis [109, 110]. In this study, we observed that the autophagy lysosomal activity for microglia and astrocytes near the implant site are comparable on day 7 post-implantation (**Fig. 4E, 4G, Fig. 6C, 6F**), indicating an orchestration of phagocytosis by microglia and astrocytes during acute implantation period. However, astrocytes significantly increased their autophagy and lysosomal activity by day 84 post-implantation, contrasting with a decreased autophagy-lysosomal activity in microglia. This may make sense as astrocytes can increase their phagocytic activity to compensate when microglial phagocytosis is genetically suppressed [106]. Thus, this increased autophagy-lysosomal activity in astrocytes suggests that astrocytes play a predominant role in phagocytosis near the chronically implanted microelectrode. Astrocytes can uptake and degrade extracellular tau via autophagy-lysosomal pathway to prevent improper accumulation of tau aggregates [111, 112]. Future research should determine the role of astrocyte clearance on tau pathologies and whether astrocytes’ phagocytic and/or autophagic activity is impaired near chronically implanted microelectrode.

### 4.3. Impaired autophagy-lysosomal activity leads to oligodendrocyte loss and myelination turnover following chronic microelectrode implantation

Oligodendrocyte soma and myelin processes are vulnerable to inflammatory injury. Instead of increasing reactivity, injury leads to loss of oligodendrocyte structural integrity and upregulation myelin-derived toxic debris such as Nogo-A and myelin-associated glycoprotein [113, 114]. These myelin debris inhibit neurite growth, axonal regeneration, oligodendrocyte precursor cell differentiation, which recruit activated microglia and reactive astrocytes to clear this debris [115, 116]. The previous study demonstrates the loss of myelin and oligodendrocyte soma starts at day 3 post implantation [30], which coincides with the peak in immature autophagy activity (**Fig. 2C**) with large, glial-like appearances (**Fig. 2F-I**). These findings suggest an acute clearance of myelin debris near the implanted microelectrode as the phagocytic activity in microglia and astrocytes can be associated with their elevated autophagy-lysosomal activity (**Fig. 6C, 6F**). This study also shows myelin debris colocalized with lysosomes in reactive astrocytes close the implant site on day 7 post-implantation (**Fig. 7**), suggesting astrocytes are clearing myelin debris through lysosomal degradation. Thus, the elevated autophagy-lysosomal activity seems to serve as a protective mechanism against the detrimental effects of injured myelin near the implanted microelectrode. However, it is possible that clearance by phagocytic cells is overwhelming and may lead to unnecessary myelin degradation, explaining the continuous demyelination injury over chronic implantation period. Thus, the elevated autophagy-lysosomal activity may act as a double-edged sword for oligodendrocyte and myelin integrity. Future investigation into the activation of autophagy-lysosomal degradation in phagocytic cells precedes myelin loss and oligodendrocyte injury near the implanted microelectrode may inspire strategies to promote myelination for neuronal functionality.

In addition, the autophagy-lysosomal-endosome axis in oligodendrocyte structures is essential for their regeneration activity [29]. Remyelination requires the expansion of the myelin membrane, which is achieved by the transport and fusion of lysosomes and endosomes containing myelin proteins [76]. Impairment of lysosomal fusion leads to abnormal myelination characterized by reduced levels of specific myelin proteins [76], highlighting the role of intracellular lysosomal activity for depositing myelin proteins for regeneration. Here, we demonstrate a transient elevation of lysosome population in oligodendrocyte soma on day 28 post-implantation near the implanted microelectrode (**Fig. 6I**), suggesting a remyelinating attempt. Interestingly, oligodendrocyte soma exhibits a significantly increased autophagy activity, yet a low level in lysosomal activity near the implant site on day 84 post-implantation (**Fig. 4I**). Oligodendrocytes use autophagy to degrade the preexisting myelin sheathes to avoid the improper myelin structures and functions [117]. Thus, the progressive sheath degradation near the chronically implanted microelectrode [30] is likely a result from the chronic elevation in oligodendrocyte autophagy for removing improper myelin structure by the implantation injury. Moreover, oligodendrocyte autophagy activity in myelin sheaths appears lower relative to activity in their soma [29]. Thus, the high colocalization of LC3B autophagy markers with MBP+ myelin at the electrode-tissue interfaces (**Fig. 4K**) likely arises from the myelinated axons. More direct evidence is required to understand whether implantation injury elevates autophagy activity in myelin and its relationship to myelin sheath loss.

### 4.4. Impaired autophagy-lysosomal activity leads to neuronal malfunctions following chronic microelectrode implantation

The chronic decline of microelectrode performance is related to the pathological features of nearby neurons, including neuronal apoptosis, axonal degeneration, and functional silencing [33, 81, 118]. Neurons, which have high metabolic demand from action potential generation, are vulnerable to inflammatory environments with high gradient of proinflammatory cytokines, oxidative stress, and cytotoxic glutamate [14, 119]. Thus, lysosome-involved degradations (e.g. autophagy, phagocytosis, endocytosis) are critical for neurons to defend against the inflammatory stressors [75]. However, we observe reduced lysosome activity in neurons adjacent to the implant over the chronic implantation period (**Fig. 5D-E**), suggesting that these neurons suffer from an impaired lysosomal clearance near the implanted microelectrode. Additionally, the increase in neurons with TFEB nuclear translocation is not accompanied by a significant increase in neuronal lysosomes microelectrode on day 84 post-implantation (**Fig, 5D-H**), suggesting a failure of lysosome biogenesis in neurons due to the chronic microelectrode implantation. Taken together, the impaired lysosomal activity may lead to the progressive neuron death and neurite degeneration reported in chronic tissue near the implanted microelectrodes.

Neurons need large amounts of energy to support action potentials and synaptic activity, and mitochondria are the major ATP-production organelles [120]. Thus, neuronal function and survival are sensitive to mitochondrial dysfunction. Mitophagy, a special form of autophagy, selectively removes damaged mitochondria [41, 121]. Previous study showed that damaged mitochondria in cultured hippocampus neurons could be quickly sequestrated in autophagosomes within an hour after initial damage [122]. However, the acidification of these immature autophagosomes takes more than 6 hours to degrade the engulfed mitochondria [122]. This slow mitochondrial turnover may be exacerbated following injury, leading to a greater buildup of damaged organelles and increasing the risk of neuronal dysfunction [122]. Here, we observe a prolonged accumulation of immature autophagy vacuoles without acidification (**Fig. 2**) together with accumulated oxidative stress (**Fig. 8G-H**), and reduced mitochondria activity (**Fig. 8E-F**) near the chronically implanted microelectrode. Taken together, chronic implantation of microelectrodes may result in inefficient removal of damaged mitochondria and thereby lead to deficits in neuron survival and action potential generation. A previous study with a long-term recording of microelectrode implanted into wild-type mice indeed demonstrates a sudden drop of neuronal detectability and SNR at week 11 post-implantation [123]. This coincidence of reduced neuronal firing and the accumulation of immature autophagy vesicles suggests that the mitophagy dysregulation in neurons may be associated with the chronic decline in device performances. Future interventions could focus on accelerating mitophagy process in chronic microelectrode implantation to prevent the loss of functional signals.

Beyond mitophagy, autophagy is vital for axonal integrity and functionality. Its deficits lead to axons swelling, retraction, and eventually degeneration [31, 124, 125]. Autophagy vesicles are predominantly found in axons relative to the soma [124], which supports our observation of the low autophagy level in NeuN+ neuronal soma at tissue-electrode interfaces (**Fig. 4C**). Autophagosomes are formed at distal axons and retrogradely move toward the soma, fusing with lysosomes to become autolysosomes during their axonal transport, ultimately acidifying in the proximal part of the axons [31, 124].

Accumulated autophagy vesicles with tau aggregates in injured axons is linked to tau pathologies in neurodegenerative diseases [50]. Microelectrode insertion disrupts the axons integrity, leading to bleb-like protrusions [118] possibly due to the disruption of microtubule structure necessary for autophagy transport and resulting in aggregation of immature autophagosomes and mature, acidified autolysosomes. Small size, dot-like morphologies of autophagy cluster near the chronically implanted microelectrode may indicate abnormal autophagy activity in injured axons. The colocalization of pathological tau proteins with demyelinated axons near the microelectrode suggests a need for investigating the role of axonal autophagy transport and abnormal hyperphosphorylated tau accumulation following microelectrode implantation.

Autophagy is also relevant to synaptic plasticity by degrading AMPA receptors and other synapse-associated proteins [126]. Similar to lysosomes, autophagosomes can fuse with late endosomes at axonal terminals, forming an intermediate organelle called amphisome [127]. Amphisomes are involved in TrkB signaling and local activation of ERK1/2 signaling at boutons for neurotransmitter release [128]. The autophagy marker LC3B can be recruited to low pH amphisome, which may express RFP-only signals [66]. Thus, the sustained RFP+ EGFP activity throughout the long-term microelectrode implantation (**Fig. 2D**) may indicate the amphisome activity in synapse remodeling at tissue-electrode interfaces.

However, distinguishing RFP signals between the lysosome-fused autolysosomes and endosome-fused amphisomes is challenging without additional techniques to examine the organelle ultrastructure. Furthermore, it is not known how the amphisomes pathway is simultaneously recruited with the autophagy pathway in the axons and their contribution to synaptic activity following the microelectrode implantation. The specific cargo identities of these pathways also remain unexplored. Further studies are needed to understand the mechanisms of protein and organelle turnover at synapses and develop interventions for promoting synaptic remodeling in neurological disorders.

### 4.5. Limitation

This study has limitations to consider. Firstly, lipofuscin autofluorescence could confound the interpretation of two-photon imaging in CAG-LC3b-RFP-EGFP in aged mice (**Fig. 1**). Lipofuscin is the complex mixture of oxidized protein and lipid residues as waste products of lysosomal digestion, which is more abundant in aged individuals [129]. Lipofuscin manifests as yellow-brown, bleb-like particles and is detectable across a broad spectrum by microscopy, which challenges the identification of yellow/green autophagosomes in the CAG-LC3b-RFP-EGFP transgenic model [57]. Therefore, to minimize the interference of age-dependent lipofuscin, 8-week young mice were used in this study. Lipofuscin-induced noise was considered negligible in 2- to 6-month-old mice [57]. However, animals were tracked for the LC3B fluorescent activity over 84-days post-implantation and were 7 months old at the end of the study. Thus, there is some possibility of lipofuscin autofluorescence especially during the last 4-week of the study. This was one of the reasons that immunohistochemistry was used to confirm LC3B findings. Characterizations of autophagosomes and lipofuscin autofluorescence clusters such as size and distribution density would help appropriately understand the two-photon imaging results.

The second limitation is the unknown origin of the chronic accumulation of immature autophagy vesicles near the chronically implanted microelectrode (**Fig. 2C**). This accumulation may be caused by increased formation of immature autophagosomes and/or impaired lysosomal fusion into RFP+ EGFP mature autolysosomes. Lysosomal fusion with autophagosomes occurs during the retrograde transport in axons [124], which means that impaired ATP-consuming motor proteins during autophagy retrograde transport may result in a decreased motility of lysosomes and autophagosomes and reduce the possibility of fusion. The dysregulation of lysosome acidification is another possible cause of the accumulated immature autophagosomes. The disruption of the blood-brain barrier following microelectrode insertion reduces metabolic supply and increases the metabolic stress [130], leading to less ATP available for the lysosome v-ATPase pump to maintain the acidic environment inside the lysosomes. In turn, this loss of ATP results in inefficient cargo degradation and chronic build-up of the immature autophagy vesicles or lipofuscin [131]. Furthermore, impairment in fusion machinery such as the interaction between Rab7A and PLEKHM1 leads to inefficient docking of lysosomes to autophagosomes [132, 133] further contributes to the accumulation of harmful autophagy cargo and in turn, exacerbates chronic implantation injury. However, the role of autophagy-lysosomal activity and the underlying mechanisms require further investigation. Understanding the chronic accumulation of autophagy vesicles may guide the development of therapeutic interventions targeting autophagosome formation and lysosomal fusion.

## 5. Conclusion

This study for the first time revealed impaired autophagy-lysosomal activity during long-term microelectrode implantation. Two photon-microscopy of CAG-RFP-EGFP-LC3 transgenic mice showed that immature autophagy vacuoles accumulate near implanted microelectrode. In addition, immunohistological results confirmed impaired autophagy-lysosomal degradation and increased oxidative stress. It is likely that reactive astrocytes contribute to myelin debris clearance and show active autophagy-lysosomal activity near the microelectrode. Overall, this study provides a novel perspective on the possible mechanisms of tissue degeneration near chronically implanted microelectrodes. These findings demonstrate that an impaired autophagy-lysosomal activity contributes to biocompatibility mechanisms of the chronic microelectrode implantation and provide insights for developing therapeutic strategies to enhance device performance.

---

**Supplementary Figure 1:**
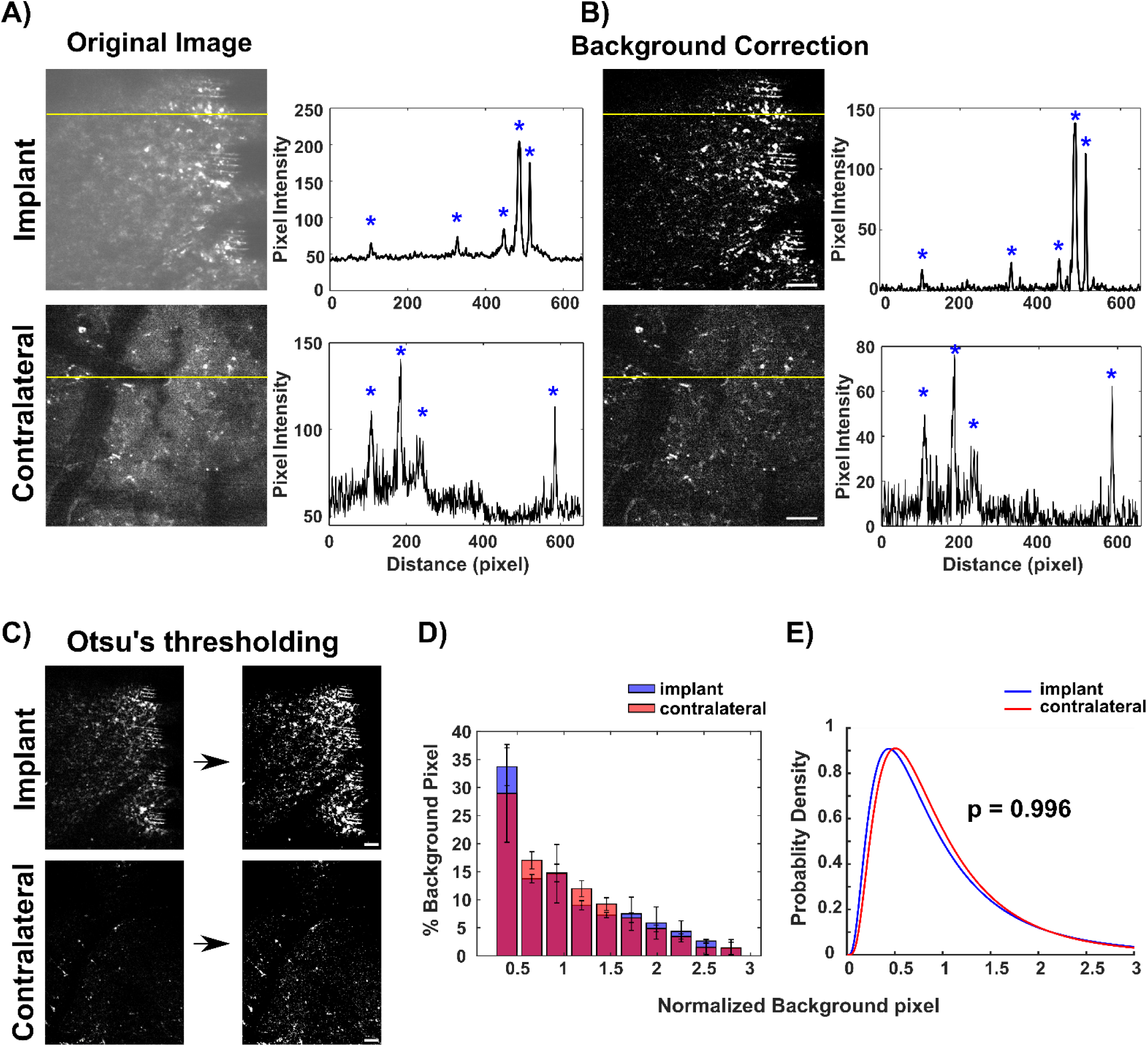
Illustration of two-photon image analysis. **A)** Left: Representative original images showing a sum z-projections of 100 μm of the visual cortex in the implant and no-implant hemispheres. Right: Profiles of fluorescence intensity across the region of interest indicated by the yellow lines. The fluorescent signals with peak amplitude are marked by blue asterisks. **B)** Left: Representative background corrected images after rolling ball background subtraction (radius = 10 pixels). Right: Profiles of fluorescent intensity within the same ROI (yellow lines). This method effectively removed most of the background gradient while preserving the features of the fluorescent signals marked by blue asterisks. **C)** Otsu’s thresholding was then applied to identify the fluorescent signals in the implant and contralateral sides. Histogram distribution **(D)** and probability density function **(E)** of normalized intensity of background pixels below Otsu’s thresholding at the implant (blue) and contralateral (red) sides. Data are presented as mean ± standard deviation. There was no significant difference in distributions of normalized intensity of background pixels between the implant and contralateral sides (Two-sample Kolmogorov–Smirnov test, *p* = 0.996). Scale bar = 50 μm.

**Supplementary Figure 2:**
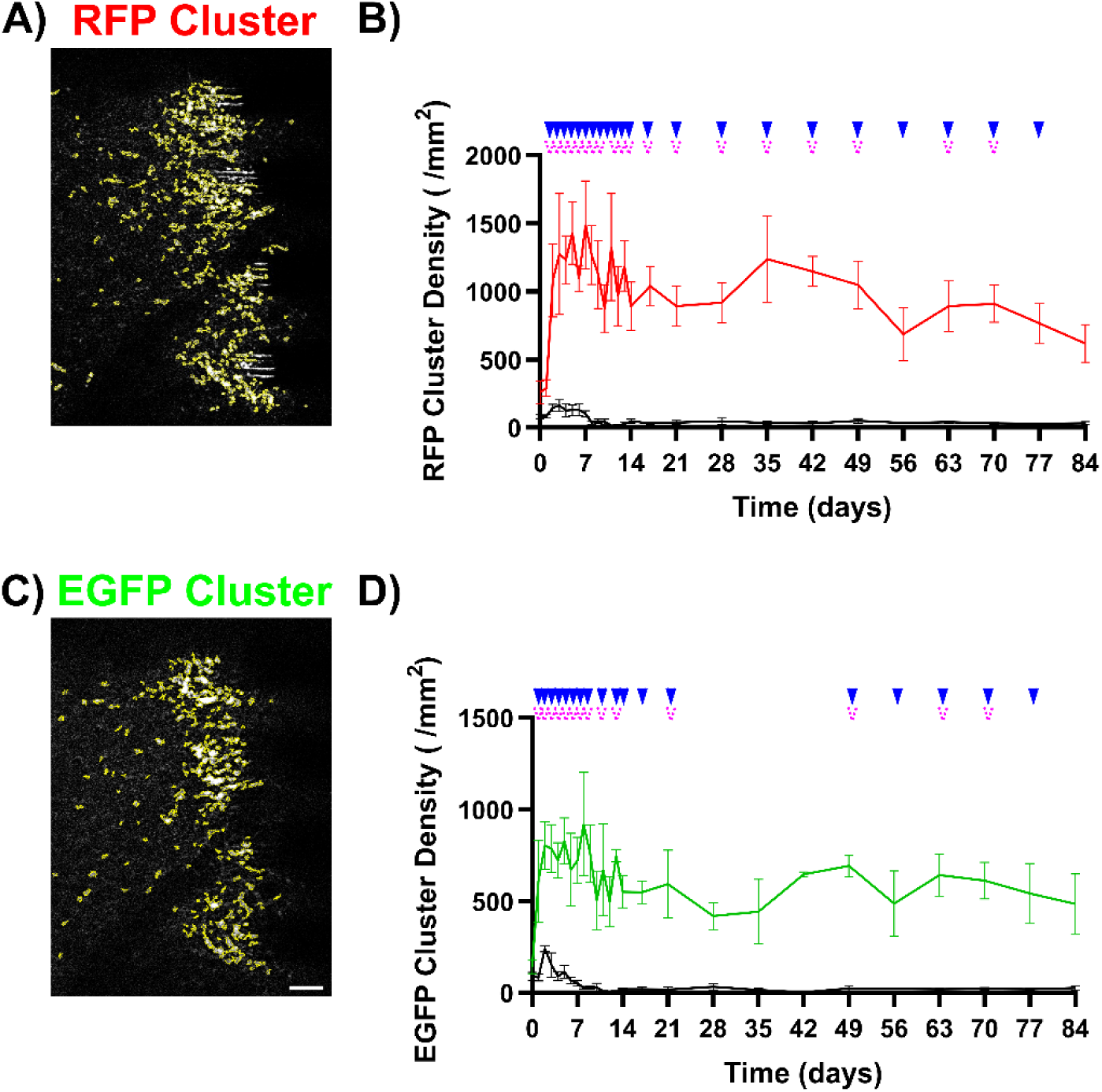
Quantification of autophagy activity based on the density of RFP and EGFP cluster near the chronically implanted microelectrode. Fluorescent signals with area greater than 1 μm^2^ (4 pixel^2^) were automatically identified as the fluorescent cluster by using a build-in ImageJ feature “Analyze Particles”. A) Representative image showing identified RFP clusters (outlined by yellow line) near the implanted microelectrode. B) Density of RFP clusters within the 250 μm x 350 μm ROI quantified at the microelectrode interfaces (colored) and no implant contralateral side (black) over time. The density of RFP clusters at the microelectrode interfaces consistently remained elevated until 84-days post-implantation. C) Representative image displaying identified EGFP clusters (outlined by yellow lines) near the implanted microelectrode. D) Density of EGFP clusters at the microelectrode interfaces (green line) exhibited a significant peak within 21-days and a significant chronic accumulation between 49-77 days post implantation compared to the contralateral side (black line). Scale bar = 50 μm. Significant comparisons between the implant and contralateral regions at each time point are denoted by blue solid triangles (*p* < 0.05), and significant comparisons between day 0 and other time points within the implant ROI are indicated by pink dash arrows (*p* < 0.05). No significant comparisons between time points at contralateral regions. All data are presented as mean ± standard deviation.

**Supplementary Figure 3:**
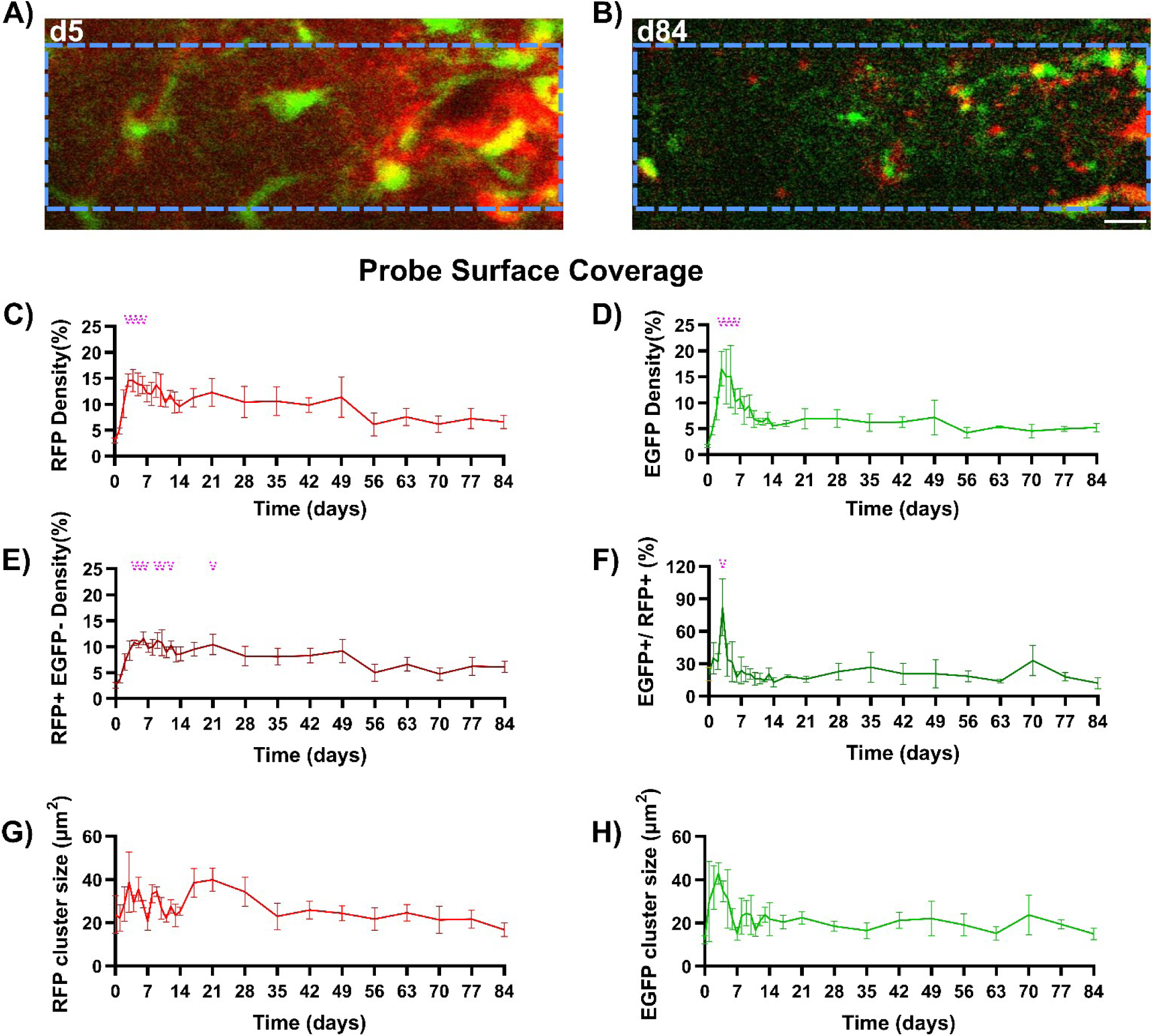
Increased autophagy activity observed over the surface of the implant during the acute inflammatory period. Representative two-photon sum projections of a small volume of tissue located 20 μm directly above the implant surface at 5- **(A)** and 84- **(B)** days post-implantation. The implant is outlined in blue dash lines. Scale bar = 20 μm. **C)** Measurement of RFP+ pixel density covering the implant surface over time, indicating a significant peak activity during days 3-6 post-implantation compared to day 0. **D)** Measurement of EGFP pixel density over the surface of the implanted microelectrode, demonstrating a significant peak activity during days 3-6 post-implantation compared to day 0. **E)** Measurement of RFP+ EGFP pixel density, representing mature autolysosome activity over the implant surface, showed a significant increase from days 4 to 21 post-implantation compared to day 0. **F)** Population ratio of EGFP pixel density over the RFP pixel density over the implant surface had a significant elevation on day 3 post-implantation compared to day 0. Changes in the size of RFP clusters **(G)** and EGFP clusters **(H)** over the implant surface during implantation period. Significant comparisons between day 0 and other time points within the implant ROI are indicated by pink dash arrows (*p* < 0.05). No significant comparisons were observed between time points in the contralateral regions. All data are presented as mean ± standard deviation.

**Supplementary Figure 4:**
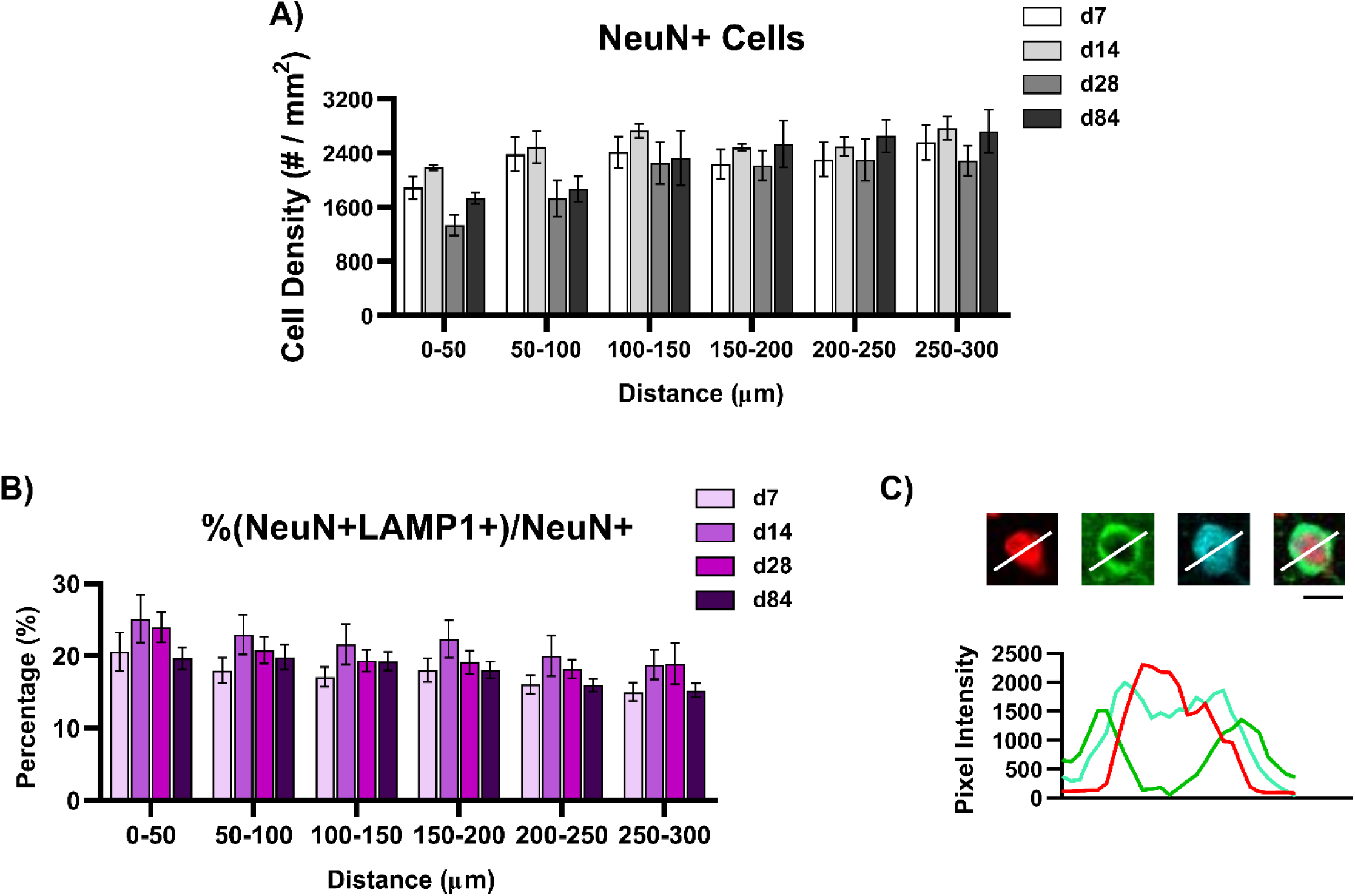
Autophagy activity in neurons near the chronically implanted microelectrode over time. **A)** Density of NeuN+ cells observed around the implanted microelectrode on days 7, 14, 28, and 84 post-implantation **B)** Percentage of NeuN pixels colocalized with LAMP1+ signals measured within 50 μm bins up to 300 μm from the implant site on days 7, 14, 28, and 84 post-implantation. **C)** Representative image showing a neuron without TFEB nuclear translocation. Scar bar = 10 μm.

**Supplementary Figure 5:**
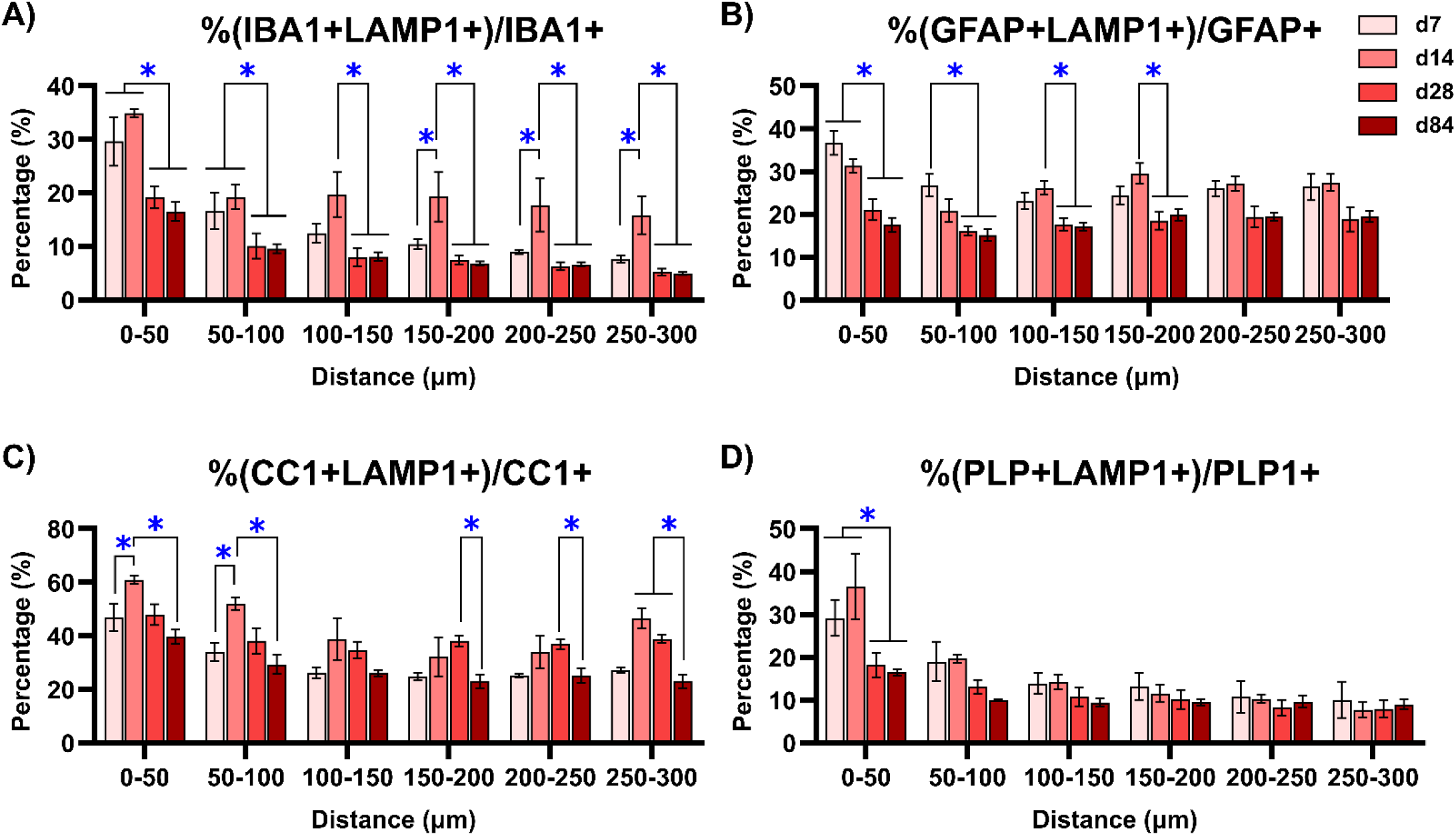
Reduced autophagy activity in glial cells near the microelectrode over the chronic implantation period. **A)** Significant reduction in the percentages of IBA1+ microglia signals colocalized with LAMP1+ signals on day 28 post-implantation over 300 μm from the implant. **B)** Significant decrease in the percentages of GFAP+ astrocyte signals colocalized with LAMP1+ markers on day 28. **C)** Significant decrease in percentages of CC1+ oligodendrocyte signals colocalized with LAMP1 on day 84 post-implantation. **D)** Significant reduction in the percentage of PLP1+ myelin with LAMP1 significantly reduced within 50 μm away from the probe on day 28 post-implantation. Blue asterisks indicate significant differences in colocalization percentage between day 7 and day 84 post-implantation at each distance bin (Two-way ANOVA with Tukey post-hoc, *p* < 0.05). All data are presented as mean ± standard deviation.

**Supplementary Figure 6:**
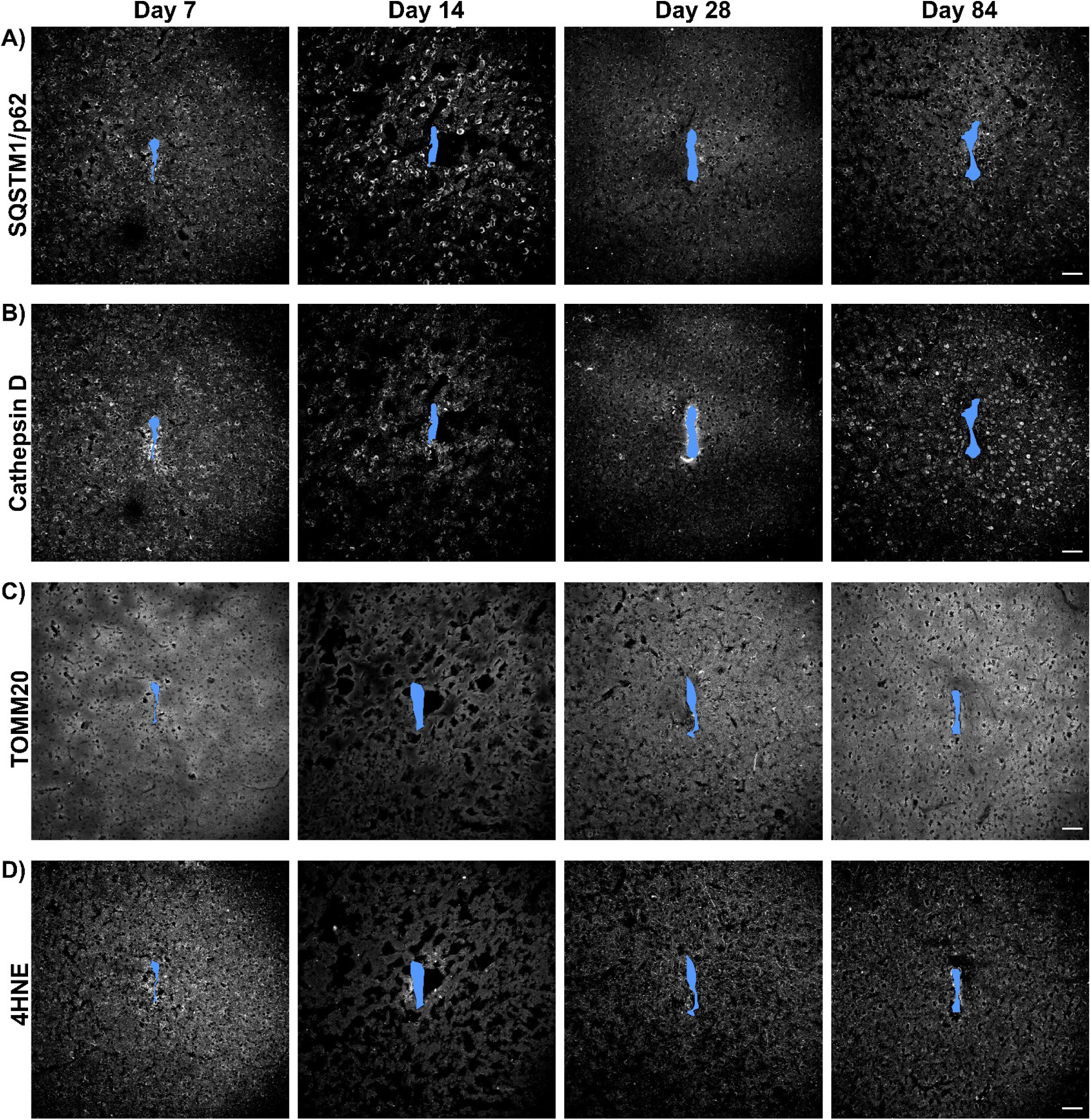
Representative immunohistological images of markers for intracellular autophagy lysosomal malfunctions at 7-, 14-, 28-, and 84-days post-implantation. Staining images of autophagy clearance marker SQSTM1/p62 (A), acidic lysosomal protease marker Cathepsin D (B), mitochondrial marker TOMM20 (C), oxidative stress marker 4-Hydroxynonenal (4-HNE) (D) near the chronically implanted microelectrode. The microelectrode implant site is marked in solid blue. Scale bar = 50 µm.

## Acknowledgment

The authors would like to thank Christopher Hughes, Naofumi Suematsu, Kevin Stieger, Fan Li for proofreading. This work was supported by NIH R01NS094396, NIH R01NS105691, NIH R01NS115707 and a diversity supplement to this parent grant, NIH R03AG072218, NSF CAREER 1943906, NIH R01NS096755,and Swanson School of Engineering (SSOE) - Department of Biological Sciences/Department of Neuroscience award funding which was jointly provided by the Senior Vice Chancellor for Research, the Dietrich School, and the Swanson School.

## Declaration of competing interest

The authors declare that they have no known competing financial interests or personal relationships that could have appeared to influence the work reported in this paper.

## Data availability

Data will be made available on request.

## Notes

http://www.bioniclab.org

### Competing Interest Statement

The authors have declared no competing interest.

