## Supplemental Figures for "Impairment of autophagy-lysosomal activity near the chronically implanted microelectrodes"

**Abstract**

Intracortical microelectrodes that can record and stimulate brain activity have become a valuable technique for basic science research and clinical applications. However, long-term implantation of these microelectrodes can lead to progressive neurodegeneration in the surrounding microenvironment, characterized by elevation in disease-associated markers. Dysregulation of autophagy-lysosomal degradation, a major intracellular waste removal process, is considered a key factor in the onset and progression of neurodegenerative diseases. It is plausible that similar dysfunctions in autophagy-lysosomal degradation contribute to tissue degeneration following implantation-induced focal brain injury, ultimately impacting recording performance. To understand how the focal, persistent brain injury caused by long-term microelectrode implantation impairs the autophagy-lysosomal pathway, we employed two-photon microscopy and immunohistology. This investigation focused on the spatiotemporal characterization of autophagy-lysosomal activity near the chronically implanted microelectrode. We observed an aberrant accumulation of immature autophagy vesicles near the microelectrode over the chronic implantation period. Additionally, we found deficits in autophagy-lysosomal clearance proximal to the chronic implant, which was associated with an accumulation of autophagy cargo and a reduction in lysosomal protease level during the chronic period. Furthermore, our evidence suggests astrocytes contribute to the clearance of myelin debris via autophagy-lysosomal degradation near the implanted microelectrode. Together, this study sheds light on the process of brain tissue degeneration caused by long-term microelectrode implantation, with a specific focus on impaired intracellular waste degradation.

**Keywords:** brain-computer interface, brain metabolism, neuron survival, glial reactivity, tissue homeostasis

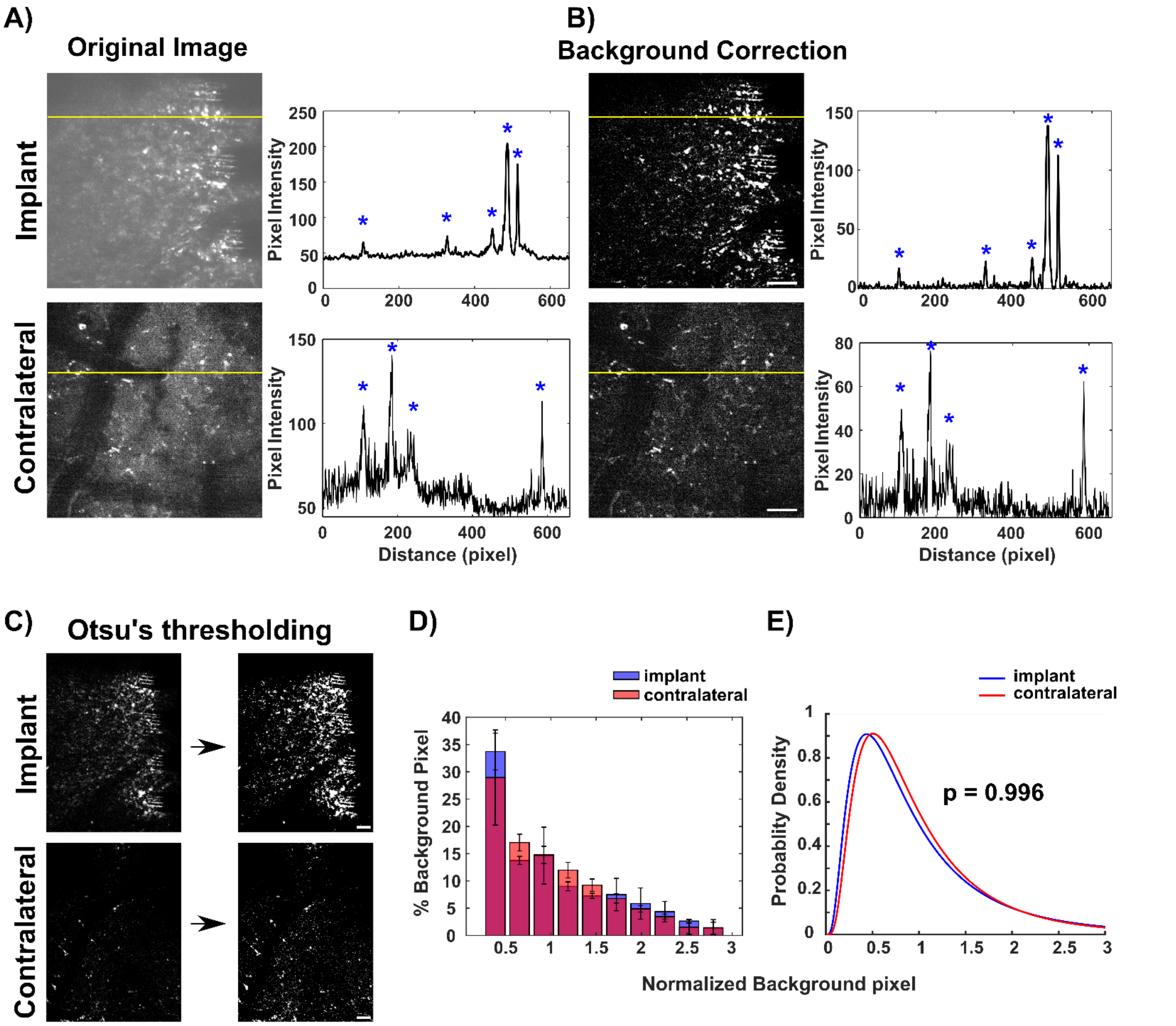


**Supplementary Figure 1: Illustration of two-photon image analysis. A)** Left: Representative original images showing a sum z-projections of 100 µm of the visual cortex in the implant and no-implant hemispheres. Right: Profiles of fluorescence intensity across the region of interest indicated by the yellow lines. The fluorescent signals with peak amplitude are marked by blue asterisks. **B)** Left: Representative background corrected images after rolling ball background subtraction (radius = 10 pixels). Right: Profiles of fluorescent intensity within the same ROI (yellow lines). This method effectively removed most of the background gradient while preserving the features of the fluorescent signals marked by blue asterisks. **C)** Otsu's thresholding was then applied to identify the fluorescent signals in the implant and contralateral sides. Histogram distribution **(D)** and probability density function **(E)** of normalized intensity of background pixels below Otsu's thresholding at the implant (blue) and contralateral (red) sides. Data are presented as mean ± standard deviation. There was no significant difference in distributions of normalized intensity of background pixels between the implant and contralateral sides (Two-sample Kolmogorov–Smirnov test, $p$ = 0.996). Scale bar = 50 µm.

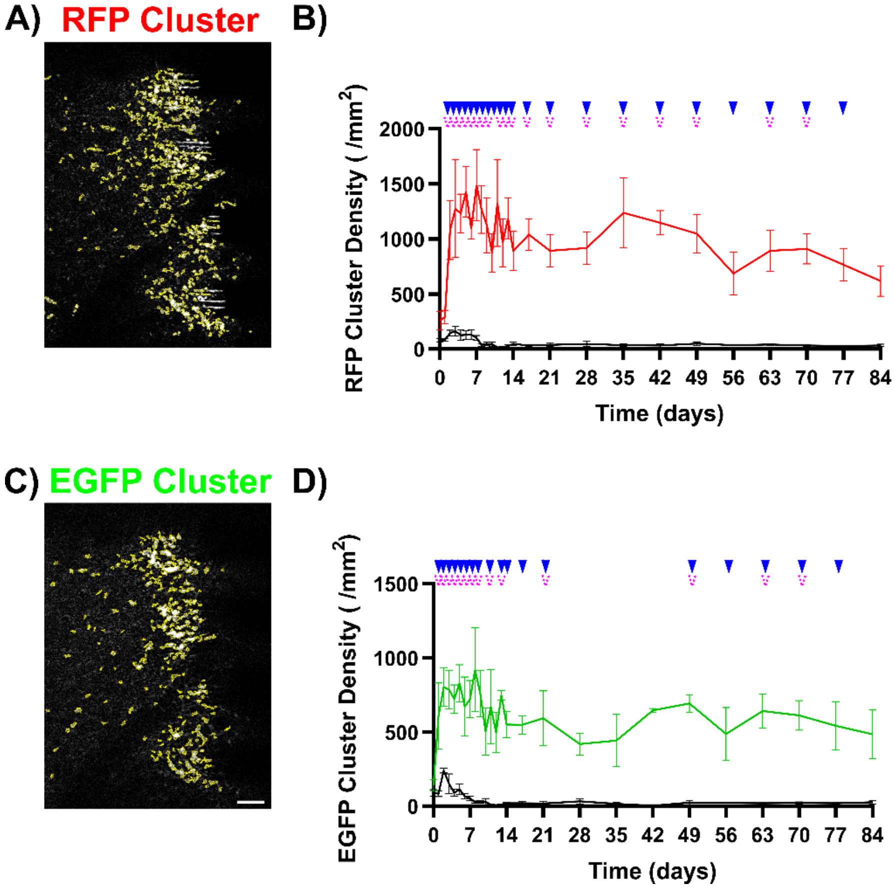


**Supplementary Figure 2: Quantification of autophagy activity based on the density of RFP and EGFP cluster near the chronically implanted microelectrode.** Fluorescent signals with area greater than 1 μm$^2$ (4 pixel$^2$) were automatically identified as the fluorescent cluster by using a build-in ImageJ feature “Analyze Particles”. **A)** Representative image showing identified RFP clusters (outlined by yellow line) near the implanted microelectrode. **B)** Density of RFP clusters within the 250 μm x 350 μm ROI quantified at the microelectrode interfaces (colored) and no-implant contralateral side (black) over time. The density of RFP clusters at the microelectrode interfaces consistently remained elevated until 84-days post-implantation. **C)** Representative image displaying identified EGFP clusters (outlined by yellow lines) near the implanted microelectrode. **D)** Density of EGFP clusters at the microelectrode interfaces (green line) exhibited a significant peak within 21-days and a significant chronic accumulation between 49-77 days post-implantation compared to the contralateral side (black line). Scale bar = 50 μm. Significant comparisons between the implant and contralateral regions at each time point are denoted by blue solid triangles ($p$ < 0.05), and significant comparisons between day 0 and other time points within the implant ROI are indicated by pink dash arrows ($p$ < 0.05). No significant comparisons between time points at contralateral regions. All data are presented as mean ± standard deviation.

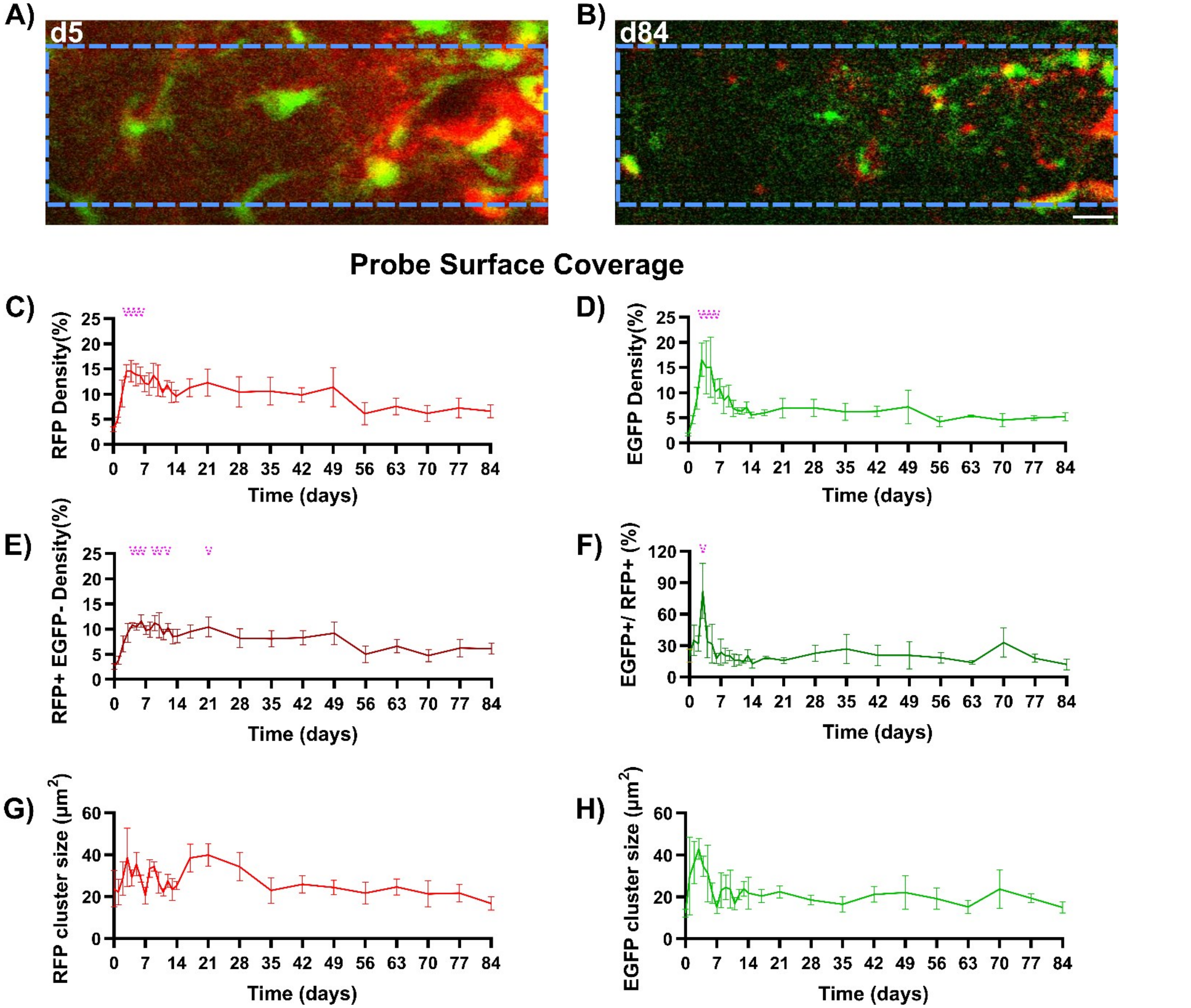


**Supplementary Figure 3: Increased autophagy activity observed over the surface of the implant during the acute inflammatory period.** Representative two-photon sum projections of a small volume of tissue located 20 μm directly above the implant surface at 5- **(A)** and 84- **(B)** days post-implantation. The implant is outlined in blue dash lines. Scale bar = 20 μm. **C)** Measurement of RFP+ pixel density covering the implant surface over time, indicating a significant peak activity during days 3-6 post-implantation compared to day 0. **D)** Measurement of EGFP pixel density over the surface of the implanted microelectrode, demonstrating a significant peak activity during days 3-6 post-implantation compared to day 0. **E)** Measurement of RFP+ EGFP- pixel density, representing mature autolysosome activity over the implant surface, showed a significant increase from days 4 to 21 post-implantation compared to day 0. **F)** Population ratio of EGFP pixel density over the RFP pixel density over the implant surface had a significant elevation on day 3 post-implantation compared to day 0. Changes in the size of RFP clusters **(G)** and EGFP clusters **(H)** over the implant surface during implantation period. Significant comparisons between day 0 and other time points within the implant ROI are indicated by pink dash arrows ($p < 0.05$). No significant comparisons were observed between time points in the contralateral regions. All data are presented as mean ± standard deviation.

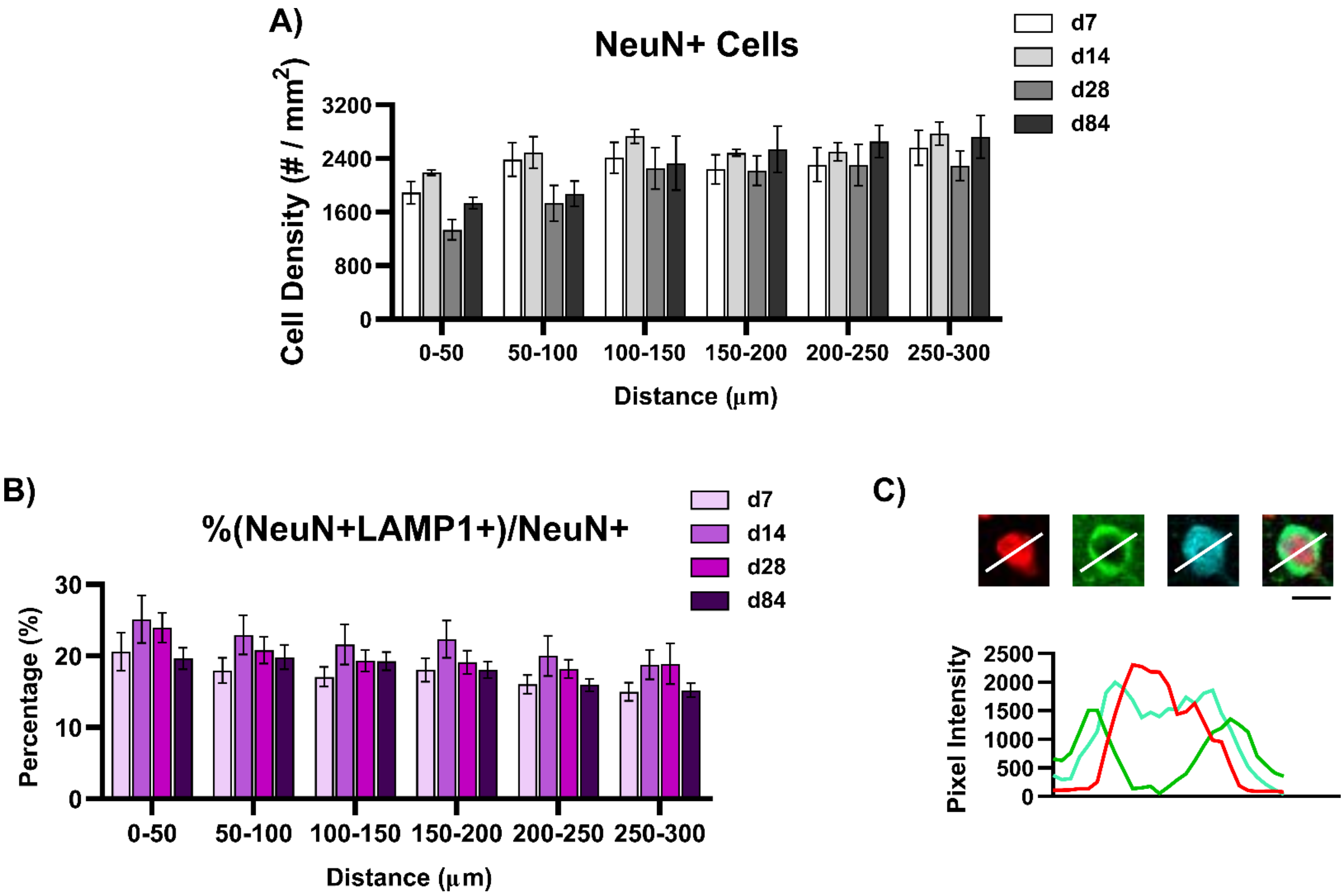


**Supplementary Figure 4: Autophagy activity in neurons near the chronically implanted microelectrode over time.** **A)** Density of NeuN+ cells observed around the implanted microelectrode on days 7, 14, 28, and 84 post-implantation **B)** Percentage of NeuN pixels colocalized with LAMP1+ signals measured within 50 μm bins up to 300 μm from the implant site on days 7, 14, 28, and 84 post-implantation. **C)** Representative image showing a neuron without TFEB nuclear translocation. Scar bar = 10 μm.

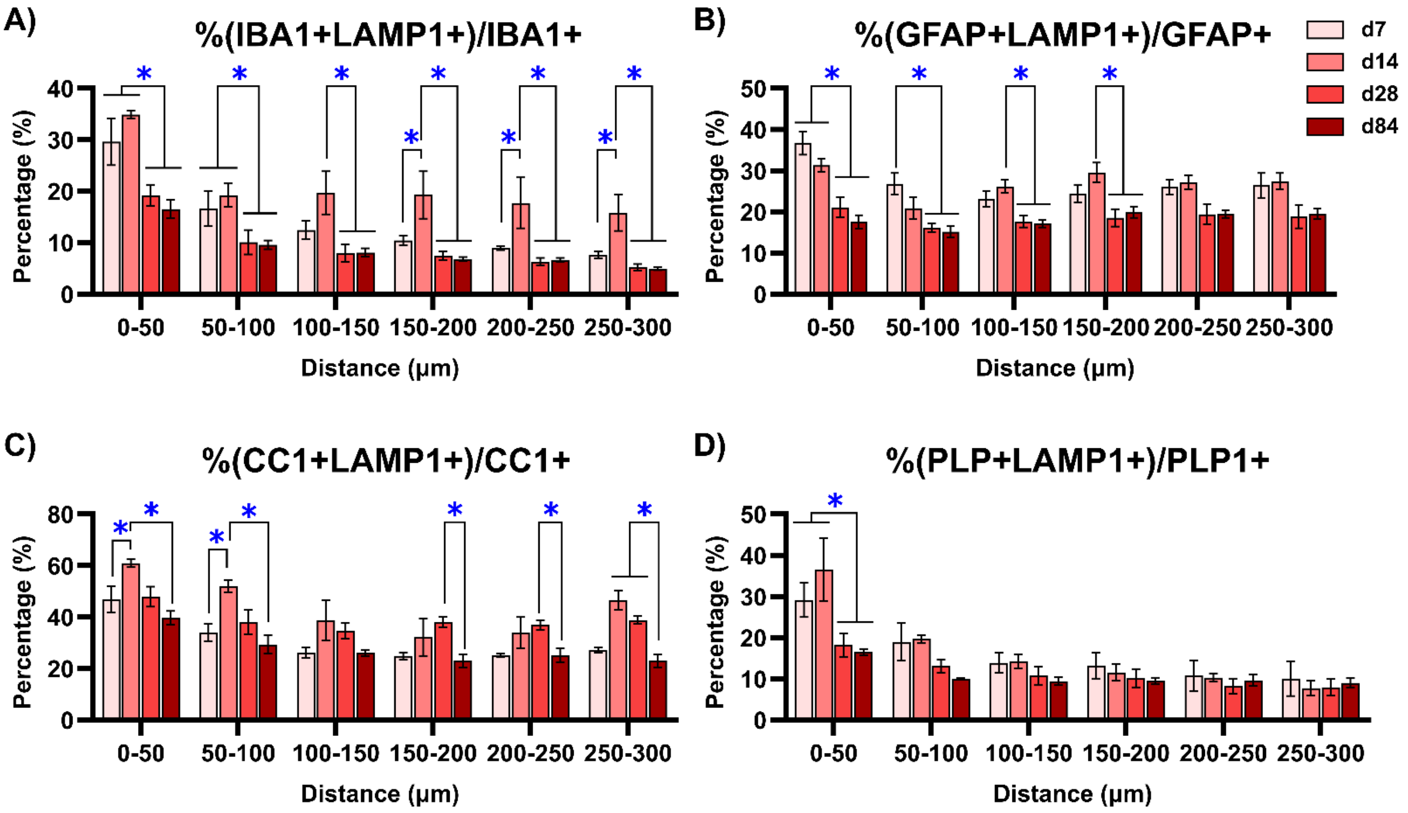


**Supplementary Figure 5: Reduced autophagy activity in glial cells near the microelectrode over the chronic implantation period. A)** Significant reduction in the percentages of IBA1+ microglia signals colocalized with LAMP1+ signals on day 28 post-implantation over 300 µm from the implant. **B)** Significant decrease in the percentages of GFAP+ astrocyte signals colocalized with LAMP1+ markers on day 28. **C)** Significant decrease in percentages of CC1+ oligodendrocyte signals colocalized with LAMP1 on day 84 post-implantation. **D)** Significant reduction in the percentage of PLP1+ myelin with LAMP1 significantly reduced within 50 µm away from the probe on day 28 post-implantation. Blue asterisks indicate significant differences in colocalization percentage between day 7 and day 84 post-implantation at each distance bin (Two-way ANOVA with Tukey post-hoc, $p < 0.05$). All data are presented as mean ± standard deviation.

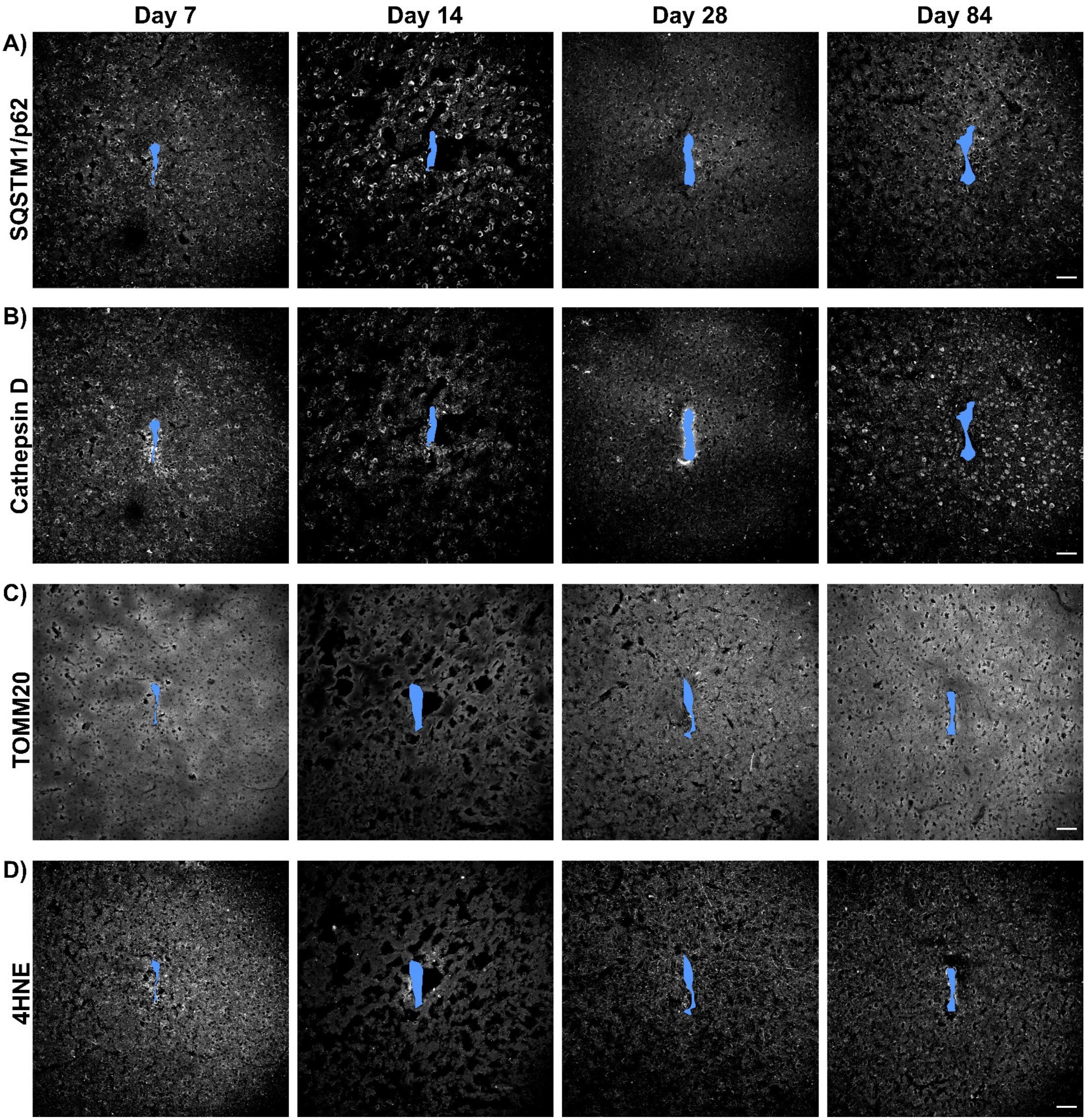


**Supplementary Figure 6: Representative immunohistological images of markers for intracellular autophagy-lysosomal malfunctions at 7-, 14-, 28-, and 84-days post-implantation**. Staining images of autophagy clearance marker SQSTM1/p62 **(A)**, acidic lysosomal protease marker Cathepsin D **(B)**, mitochondrial marker TOMM20 **(C)**, oxidative stress marker 4-Hydroxynonenal (4-HNE) **(D)** near the chronically implanted microelectrode. The microelectrode implant site is marked in solid blue. Scale bar = 50 µm.
